# Phenotypic and genotypic parallel evolution in parapatric ecotypes of *Senecio*

**DOI:** 10.1101/2020.02.05.936450

**Authors:** Maddie E. James, Melanie J. Wilkinson, Diana M. Bernal, Huanle Liu, Henry L. North, Jan Engelstädter, Daniel Ortiz-Barrientos

## Abstract

The independent and repeated adaptation of populations to similar environments often results in the evolution of similar forms. This phenomenon creates a strong correlation between phenotype and environment and is referred to as parallel evolution. However, we are still largely unaware of the dynamics of parallel evolution, as well as the interplay between phenotype and genotype within natural systems. Here, we examined phenotypic and genotypic parallel evolution in multiple parapatric Dune-Headland coastal ecotypes of an Australian wildflower, *Senecio lautus*. We observed a clear trait-environment association within the system, with all replicate populations having evolved along the same phenotypic evolutionary trajectory. Similar phenotypes have arisen via mutational changes occurring in different genes, although many share the same biological functions. Our results shed light on how replicated adaptation manifests at the phenotypic and genotypic levels within populations, and highlights *S. lautus* as one of the most striking cases of phenotypic parallel evolution in nature.

## Introduction

When separate populations are faced with similar selective pressures, they often evolve similar phenotypes (Schluter 2000). When these independent populations evolve from similar initial conditions, this phenomenon is referred to as ‘parallel evolution’ (Schluter and Nagel 1995). The correlation that arises between phenotype and environment during parallel evolution provides strong evidence for the role of natural selection in creating new forms. This is because it is unlikely that similar phenotypes would have evolved multiple times purely by chance (Lenormand et al. 2009, but see Losos 2011). Systems of parallel evolution are unique as they provide natural replicates of the evolutionary process, enabling researchers to examine the genetic architectures that modulate repeatability and determinism in nature. Although parallel evolution has been observed in a variety of animals (e.g., Nosil et al. 2002; Colosimo et al. 2005; Elmer et al. 2010; Butlin et al. 2014; Soria-Carrasco et al. 2014) and in some plants (e.g., Foster et al. 2007; Trucchi et al. 2017; Cai et al. 2019; Konečná et al. 2019; Knotek et al. 2020; Bohutínská et al. 2021), we are still largely ignorant of how the repeated adaptation to similar environments manifests at the level of the phenotype and genotype across empirical systems.

Researchers of phenotypic parallelism traditionally ask to what extent replicate populations adapted to similar environments (collectively referred to as an ‘ecotype’) are phenotypically distinct from other such ecotypes, and which traits contribute to these differences (see Bolnick et al. (2018) for a detailed review of approaches). Yet, even in some of the most classic cases of parallel evolution, such as the threespine stickleback, there can be a large degree of within-ecotype phenotypic variability between populations (Stuart et al. 2017). This has prompted a recent shift to multivariate geometric approaches (see Collyer and Adams 2007; Adams and Collyer 2009; Collyer et al. 2015; Bolnick et al. 2018; De Lisle and Bolnick 2020), which quantify how evolution proceeds within multivariate trait space and how this differs between pairs of contrasting ecotypes (as undertaken in Elmer et al., 2014; Kusche et al., 2015; Oke et al., 2017; Stuart et al., 2017; Paccard et al., 2019; Pilakouta et al., 2019; Jacobs et al., 2020). The phenotypic heterogeneity observed within natural systems highlights that evolution does not necessarily favor the exact same phenotypic features during replicated adaptation. This may be driven by a number of forces including the demographic history of populations, within-habitat environmental variation, the relationship of the phenotype to fitness landscapes, and is likely highly dependent on the underlying genetic architecture of adaptive traits (see Rosenblum et al. 2014; Lenormand et al. 2016; Blount et al. 2018 for reviews).

At the genetic level, similar phenotypes within the same environment can evolve via independent and repeated selection on the same nucleotide site or gene (reviewed in Wood et al. 2005; Christin et al. 2010; Stern 2013). For instance, replicate populations of threespine stickleback show reduction of pelvic armor due to repeated selection of alleles within the *Eda* gene (Colosimo et al. 2005). Similar phenotypes can also arise by selection on entirely different alleles and genes, although often from the same functional pathway (e.g., the parallel evolution of red flowers in *Iochroma*; Smith and Rausher 2011). In these cases, different genetic routes can produce similar phenotypic outcomes across populations, suggesting that evolution can be somewhat flexible and redundant at the level of the allele or gene. This may be especially common in systems of polygenic adaptation, where many alleles of small effect contribute additively to the adaptive phenotype (Chevin et al. 2010; Yeaman 2015; Barghi et al. 2020). Understanding the dynamics of parallel evolution will therefore allow us to gain insight into the interplay between phenotype and genotype, and will further shed light on the levels of organization at which evolution is repeatable and predictable within nature (Stern and Orgogozo 2009; Blount et al. 2018).

We must note that in the literature, the term *parallel evolution* or *parallelism* has been used quite fluidly to refer to different components of parallel evolution, including the phenotype and/or the genotype. It is perhaps not surprising that there has been a longstanding debate on the use of the term ‘*parallel evolution*’ when describing natural systems (see Haas and Simpson 1946; Arendt and Reznick 2008; Stern 2013; Lenormand et al. 2016; Stuart 2019). To reduce confusion, we hereafter avoid using ‘*parallel evolution*’ in isolation and are explicit when referring to patterns of replication that arise either at the phenotypic or genotypic levels (as suggested by Elmer and Meyer 2011). We also acknowledge that genotypic parallelism encompasses different levels of biological organization (the nucleotide site, gene, or biological function).

Here, we examine the extent of phenotypic and genotypic parallel evolution within an Australian wildflower species complex, *Senecio lautus*. The *Senecio lautus* species complex contains a variety of ecotypes adapted to contrasting environments. The Dune and Headland ecotypes are of particular interest as they consist of multiple parapatric Dune-Headland population pairs along the Australian coastline (Figure 1A) that are often sister groups in the phylogeny (Figure 1B; Roda et al. 2013a; Melo et al. 2019; James et al. 2021). Despite the close geographic proximity between populations of a pair (i.e., ecotypes within each locality), there is little to no gene flow between populations within each locality, as well as between populations within each ecotype (James et al. 2021). Previous coalescent modelling suggests that these low levels of gene flow are not high enough to create a false picture of parallel evolution, suggesting a large number of independent and repeated origins within the system (see James et al. 2021 for details). There is a strong association between overall morphology and habitat in this coastal system: Dune plants, colonizing the sandy dunes, are erect with few branches, whereas Headland individuals grow on rocky headlands and are prostrate with many branches (Figure 1C; Walter et al. 2018a). Populations maintain their phenotypes when grown in common garden conditions (Walter et al. 2016, 2018a; Wilkinson et al. 2021), suggesting that phenotypic plasticity within the system is weak. Previous work with *S. lautus* in common garden conditions has identified a suite of divergent traits between Dune and Headland populations, which include characteristics related to plant architecture and leaf morphology (Walter et al. 2018a). However, we lack a comprehensive characterization of how parallel the phenotypes and genotypes are within *S. lautus* natural populations, and how this affects divergence at the level of the ecotype and across replicate populations.

**Figure 1.**
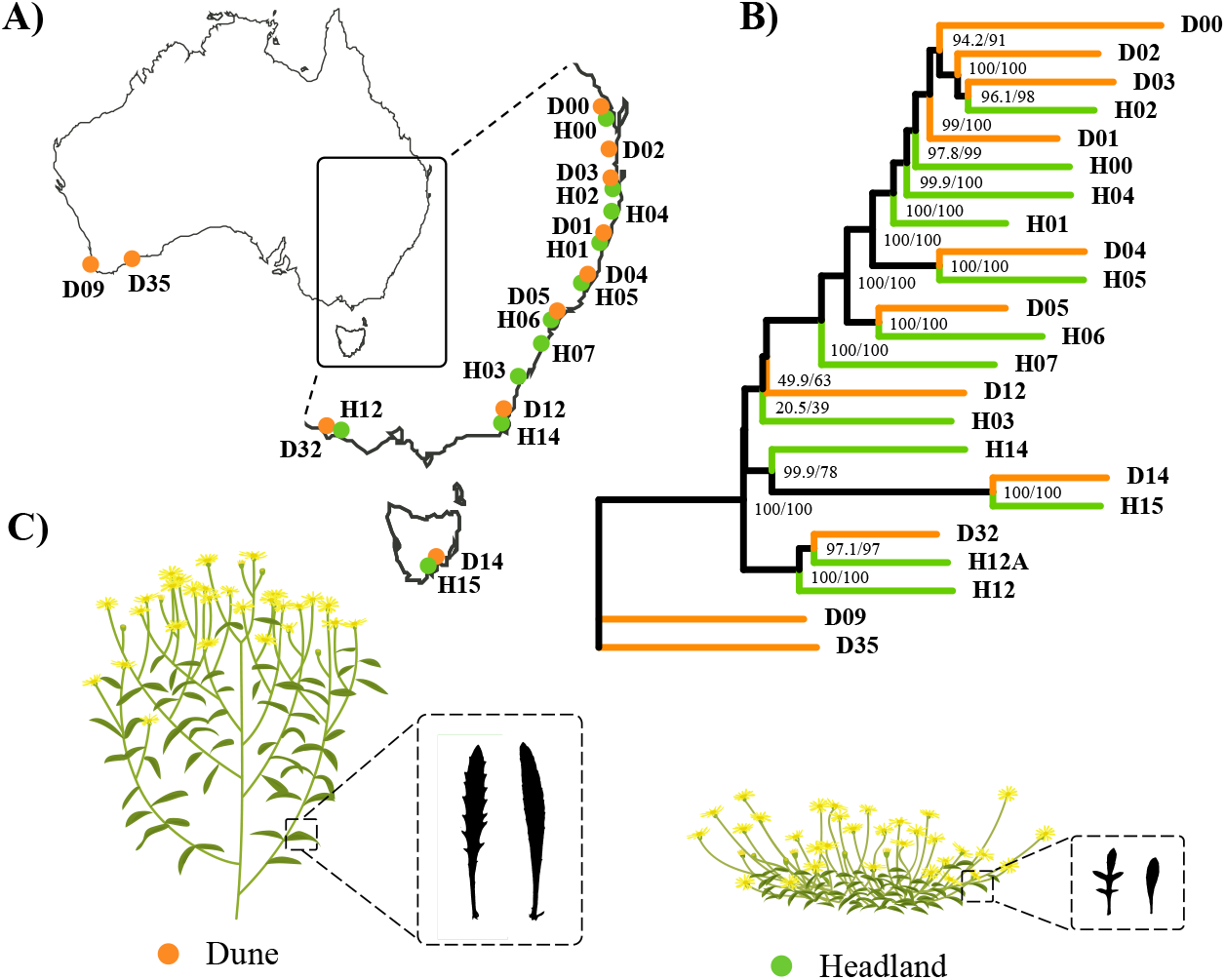
*Senecio lautus* distribution, phylogeny and ecotypes. **(A)** Sampling locations of the 22 Dune (orange) and Headland (green) *Senecio lautus* populations along the coast of Australia. **(B)** Maximum likelihood phylogeny of Dune and Headland populations implemented in IQ-TREE. Numbers on each node represent the SH-alRT support (%), followed by the ultrafast bootstrap support (%). Modified with permission from James et al. (2021). Population H12A is not included within this study. **(C)** Schematic diagram of Dune and Headland ecotypes based on mean trait values from linear discriminant analysis (LDA) shown in Figure 2A.

To assess the extent of phenotypic and genotypic parallelism in *S. lautus*, we use nine replicate Dune-Headland population pairs, two allopatric Dune populations, and two allopatric Headland populations, for a total of 22 populations. We first quantify how phenotypically distinct the Dune and Headland ecotypes are, and how this varies across the replicate population pairs at each locality. We then ask whether similar genetic mechanisms underlie these repeated phenotypes (i.e., repeated selection on the same nucleotide site, gene, or biological function), which builds on previous work using pooled sequencing of six natural Dune-Headland pairs (Roda et al. 2013b). In addition, we ask whether the variation in the extent of parallelism can be attributed to non-stochastic factors, including levels of gene flow and within ecotypic environmental variation. Overall, our work sheds light on the dynamics of parallel evolution within plants, and highlights that strikingly similar phenotypes can repeatedly evolve via different genetic routes.

## Methods

### Phenotypic parallelism

To quantify the extent of phenotypic parallelism within *S. lautus*, we measured a suite of plant architecture and leaf morphology traits from 20 Dune and Headland populations along the coast of Australia (n_mean_ = 30 individuals, n_total_ = 605; Figures 1A and 1B; Table S1). These populations include nine Dune-Headland pairs (eight which are parapatric, of which five are sister taxa; Roda et al. 2013b; Melo et al. 2019; James et al. 2021), as well as two allopatric populations that do not belong to a pair. Population pairs are based upon their geographic distribution, where a pair consists of a Dune and Headland population that are closest geographically (i.e., at the same locality). We note this is the case for all pairs, except for population D02 that we paired with H04, which are not geographic neighbors, but both reside within the eastern clade. Each *S. lautus* natural population occupies a distinct geographic range. We sampled mature (flowering) plants evenly across the range of each population, ensuring that each plant was more than one meter apart. We measured six plant architecture traits (vegetative height, widest width, narrowest width, main stem angle, main stem diameter, and primary branch angle) and eight leaf traits (area, perimeter, width, height, elongation, compactness, dissection and circularity; defined in Table S2). All plant architectural traits were measured in the field, and we sampled three primary branch leaves per plant for leaf morphometric analysis in ImageJ v1.51 (Schneider et al. 2012). Leaves were scanned at 600dpi on a CanoScan 9000F scanner and ImageJ was used to automatically extract leaf shape characteristics. Overall, these phenotypes in the wild are highly correlated with those measured under controlled conditions (Wilkinson et al. 2021).

In ~11% of individuals, we were unable to measure all six plant architectural traits (such as main stem diameter and main stem angle). In these cases, we took the average of the population to impute the trait value for that individual. We ran the below analyses with and without these individuals and obtained consistent results. We report the analyses undertaken using the population means for the missing data. All phenotypic analyses were undertaken in R v3.4.2 (R Core Team 2017). Traits were log transformed and standardized to have a mean of 0 and standard deviation of 1. We calculated pairwise correlations between all traits and removed five traits with high correlations across all populations (such that our final set of traits contained correlations < 0.8; Table S2), leaving nine traits total. These correlated traits added minimal additional phenotypic information and are thus effectively redundant.

To investigate whether the Dune and Headland ecotypes are phenotypically distinct within multivariate space, we performed a one-way MANOVA (*traits = ecotype*) across the 20 Dune and Headland populations, where the term *traits* denotes the multivariate response variable of all traits, and *ecotype* is a fixed effect of Dune or Headland. We also split traits into a plant architectural and a leaf trait-set to ask whether phenotypic differences between ecotypes depend on the trait category. Using all traits, we also performed a two-way MANOVA including *pair* as a fixed effect (*traits = ecotype + pair + ecotype × pair*), and calculated Wilk’s partial η^2^ (Langerhans and DeWitt 2004) for each term in the model using the *etasq* function in the *heplots* package (Fox et al. 2018) in R. As this model requires population pairs, we excluded two Headland allopatric populations (H03 and H07) and two Dune allopatric populations (D09 and D35). Within the two-way MANOVA, the partial effect size of the *ecotype* term denotes how much of the phenotypic variation is explained by the overall differences between ecotypes (parallel evolution), whereas the *pair* and *interaction* terms indicate how much variation is unique to replicate pairs (non-parallel evolution).

To ask whether we can predict the ecotype each individual belongs to, based on their multivariate phenotype, we performed K-means clustering with the Hartigan-Wong algorithm (Hartigan and Wong 1979). We used 25 random initial configurations and retained the run with the smallest sums of squares of the individuals to their assigned cluster center, and then calculated the proportion of individuals assigned to their correct ecotype. We also performed a linear discriminant analysis across all traits to ask which linear trait combination best explains the phenotypic differences between Dune and Headland ecotypes.

We further explored the phenotypic differences between the Dune and Headland ecotypes at a univariate trait level. We first undertook vote-counting by calculating the mean trait value for the Dune and Headland of each replicate pair and asking whether there was a consistent increase or decrease in the trait value for all replicate pairs (two-sided dependent-samples sign-tests). As this approach requires population pairs, we again excluded the four allopatric populations (H03, H07, D09 and D35). However, this vote-counting approach ignores trait effect-size, and has low statistical power when the sample size (number of replicate pairs) is small. Therefore, we used trait-by-trait linear models (ANOVAs: *trait = ecotype + pair + ecotype × pair*) to ask whether there was a significant main effect of ecotype for each trait. We also extracted the partial effect sizes (partial η^2^; Langerhans and DeWitt 2004) for each term in the model using the *etasq* function in the *heplots* package (Fox et al. 2018) in R.

We quantified the direction and magnitude of phenotypic divergence of each replicate Dune-Headland population pair using Phenotypic Change Vector Analysis (PCVA; Collyer and Adams 2007; Adams and Collyer 2009; Collyer et al. 2015). Within multivariate phenotypic space, PCVA quantifies both 1) the magnitude of divergence, and 2) the contribution of traits to divergence between replicate pairs. The procedure is as follows: the phenotypic centroid (multivariate mean) is calculated per population. For each population pair (i.e., the Dune and Headland at each locality), their centroids are connected with a vector. The length (L) of this vector quantifies how divergent the two populations are – the greater the length, the more divergent. The difference in length (ΔL) between vectors thus denotes the difference in the magnitude of divergence between two replicate population pairs. The two pairs are considered parallel with regards to the magnitude of their divergence if ΔL is not statistically different from zero (ΔL ≈ 0; Bolnick et al. 2018).

The contribution of traits to divergence is measured by the angle between vectors (θ). A large angle between two pairs (θ ≫ 0°) suggests the traits contributing to population divergence are quite different between the pairs. The contribution of traits is considered parallel when the angle is not statistically different from zero (θ ≈ 0°). Using R code modified from Collyer and Adams (2007), we calculated ΔL and θ for all pairwise comparisons between localities and performed permutations to test for statistical significance. To ensure this analysis was robust and not dominated by a single trait, we repeated the calculations of ΔL and θ nine times, removing a single trait each time. We observed consistent results across all calculations, suggesting our results are not dominated by a single trait (Table S3).

Although the above pairwise comparisons can inform us about whether the phenotypic divergence between ecotypes is similar across pairwise localities, it does not adequately assess whether evolutionary change has been more parallel than expected by chance (De Lisle and Bolnick 2020). For instance, many pairwise angles may be statistically different from zero (θ ≈ 0°; i.e., non-parallel), yet the divergence between ecotypes across localities may share a common axis of evolutionary change (De Lisle and Bolnick 2020). These common axes of divergence are not captured by PCVA, so interpreting the individual pairwise comparisons between localities can give a false impression of the extent of phenotypic parallel evolution. We therefore used a complementary approach by De Lisle and Bolnick (2020) to identify the major axes of shared evolutionary change across replicate populations, and to assess the extent of multivariate parallel evolution in the system. More specifically, we used a modified approach from De Lisle and Bolnick (2020) to calculate the correlation matrix of the matrix of individual pairwise angles between Dune-Headland replicate populations at each locality (after first normalizing the angles to radians). We then used eigenanalysis to calculate the major axes of shared evolutionary change. To identify significant axes, we generated a null distribution by sampling from an eight-dimensional Wishart distribution with nine degrees of freedom. The null expectation of no shared axes of evolutionary change was represented by an identity matrix. For each eigenvector, we sampled from this distribution 100 times. We then calculated the strength of parallelism, which is the proportion of variance that is explained by the significant eigenvectors (as identified above). We examined the loadings of the significant eigenvectors to understand how each of the population pairs contribute to parallel evolution.

### Genotypic parallelism

To quantify the extent of genotypic parallelism within *S. lautus*, we used nine Dune-Headland population pairs from a previous Genotyping-by-Sequencing dataset generated from James et al. (2021) (n_mean_ = 56 individuals, n_total_ = 1009; Figures 1A and 1B; Table S1). See James et al. (2021) for details on DNA extraction, library preparation and bioinformatics. We filtered for an overall minor allele count of five, retaining 9,269 single nucleotide polymorphisms (SNPs) across all populations. We note that linkage disequilibrium decays quickly in the system, where the mean size of a haploblock is 359bp, and the median is 42bp. Therefore, the SNPs we sampled can be largely treated as independent (see Figure S1 for more details).

#### Identifying parallel nucleotide polymorphisms

We first characterized how much genotypic variation of each of the 9,269 sequenced SNPs is explained by the overall differences between ecotypes compared to the individual replicate pairs at each locality. More specifically, we used PLINK v1.9 (Purcell et al. 2007) to normalize each SNP by conducting a PCA and extracting the loadings of the first eigenvector across all individuals. For each SNP we used these loadings to perform linear models in R (ANOVA: *SNP = ecotype + pair + ecotype × pair*) and extracted the partial effect sizes (partial η^2^) for each term in the model. To plot these data as a frequency distribution, we calculated each SNP’s distance from a 1:1 line by subtracting the effect size for either the *pair* or *interaction* term from the *ecotype* term in the model. Positive values indicate more parallel evolution (as the Dune-Headland evolutionary divergence is shared across replicate localities), whereas negative values indicate more non-parallel evolution, as the divergence is more unique to individual replicate localities.

We further explored the detailed patterns of parallelism at the level of the nucleotide site by undertaking three complementary approaches. *Approach 1*: we detected overall outliers comparing all Dune populations vs all Headland populations (using a combination of top *F_ST_* values, top cluster separation scores (CSS; Jones et al. 2012) and *BayeScan* (Foll and Gaggiotti 2008). *Approach 2*: we detected outliers separately for the Dune-Headland pairs at each locality (again using a combination of top *F_ST_* values, top CSS and *BayeScan*) and asked which SNPs were shared outliers in multiple replicate pairs. We also calculated the number of shared outlier SNPs between all pairwise comparisons across localities, and asked whether the number of shared outliers was greater than expected by chance by using a hypergeometric distribution function, *phyper*, within R. *Approach 3*: if a nucleotide site was detected as highly differentiated in at least one pair from *Approach 2*, we compared allele frequencies across all pairs for the site, and we asked whether the Δ*p* for each replicate pair was in the same direction across all nine or eight localities. Our overall best candidates SNPs for parallelism at the nucleotide site are loci that overlap between the three methods, i.e., they show high differentiation between ecotypes (*Approach 1*), high differentiation within each replicate pair (*Approach 2*) and have concordant allele frequency changes across replicate pairs (*Approach 3*). See Supplementary Methods S1 for the specific details of each approach.

To ask whether the candidate outliers from any of the approaches above fall within genic or non-genic regions, we used the first version of the *S. lautus* transcriptome (Liu 2014) (See Supplementary Methods S2). We mapped the transcriptome to the reference PacBio genome v1.0 (James et al. 2021) with *minimap2* v2.17 (Li 2018) using default parameters. We considered each transcript a separate gene, which included all isoforms. As the transcriptome excludes introns, we still considered SNPs mapped to the reference genome that fall between two segments of the same transcript as a genic SNP. All other SNPs were considered non-genic, which are expected to include variants in regulatory and repetitive regions as well as in genic regions with unknown homologous genes in other plants. We excluded SNPs that had > 1 gene mapping to it.

#### Identifying parallel genic polymorphisms

As with the nucleotide sites, we assessed the extent to which genic variation captured with protein-coding sites is explained by the differences between ecotypes compared to the individual replicate pairs at each locality. We again normalized the data, retaining the loadings of the first eigenvector for each gene. For each gene we performed linear models (ANOVA: *gene = ecotype + pair + ecotype × pair*) and extracted the partial effect sizes (partial η^2^) for each term in the model. We plotted this as a frequency distribution (see above for details).

For the SNPs detected above as overall best candidates for the SNP parallelism, we explored how many genes they fall in, and what their functions are. We also did this for SNPs detected as outliers in *Approach 1* and *Approach 3* separately. We note that we do not consider *Approach 2* as we did not detect any outliers across all nine or eight replicate pairs in *Approach 2* (see *Results* below). To assign orthologous genes, we obtained a RefSeq code per gene (Pruitt 2004) by using *BLASTx* (Altschul et al. 1990) with the *S. lautus* transcript in which the outlier SNP fell within. We searched the RefSeq protein database for *Arabidopsis thaliana* proteins that match our target genes using an E-value threshold of < 10^−6^. We used the web-based version of *DAVID* v6.8 (Huang et al. 2009b,a) to obtain the predicted functional annotation of each *S. lautus* gene sequenced in this work.

We further examined patterns of gene parallel evolution between pairwise localities by calculating the number of shared outlier genes between all pairwise comparisons across localities, and assessed whether the number of shared genes were greater than expected by chance by using a hypergeometric distribution function, *phyper*, within R. We considered a gene an outlier per replicate pair if it harbored at least one differentiated SNP according to *Approach 2* above.

#### Identifying enriched biological functions

To understand whether the outliers per population pair were enriched for any functional categories, we conducted a gene-enrichment analysis for the outlier genes for each replicate pair using functional annotation clustering in *DAVID*, using the *Arabidopsis* orthologues for our outlier genes. Functional annotation clustering groups similar functional terms into clusters to avoid redundant annotations. We considered a cluster as enriched if at least one category within the cluster had a P-value < 0.05 (the EASE score, calculated using a modification of Fisher’s exact test; Huang et al. 2009b,a). See Supplementary Methods S3 for details. We compared these enriched clusters across localities to ask whether any biological functions were repeatedly enriched across the entire system. We then used a two-sided dependent-samples sign-test to ask if the number of enriched pairs per predicted functional category differed from chance. For these enrichment analyses, the *Arabidopsis thaliana* genome was used as a genetic background, as done with previous work within *S. lautus* (Roda et al. 2013b; Wilkinson et al. 2021). We currently lack an annotated reference genome, precluding us from using *S. lautus* as a reference genome. Finally, we compared the distributions of the proportions of shared outlier nucleotide sites, outlier genes and enriched biological functions across pairs using a two-sided *X*^2^-test with continuity correction in R using the *prop.test* function.

### Demographic effects on phenotypic parallelism

Next, we tested whether the variation in phenotypic parallelism within the system (i.e., differences in divergence (ΔL) and the contribution of traits (θ) between replicate pairs) could be explained by demographic factors. We used gene flow estimates from James et al. (2021) (which were estimated from the same dataset as used within this study) to ask whether gene flow constrains divergence (linear model: *phenotypic length (L) = gene flow*; Table S4). We also used divergence time estimates from James et al. (2021) to ask whether older pairs show more phenotypic divergence than younger pairs as they have experienced more genetic drift over time (linear model: *phenotypic length (L) = divergence time*; Table S4). We also reasoned that populations adapting to more contrasting environments should have greater phenotypic differences (linear model: *phenotypic length (L) = environmental distance*; Table S4). We used environmental distances from previous work in *S. lautus* (see Roda et al. 2013b), which consisted of 38 variables of soil composition of the natural populations. In addition, we asked whether pairs that were more phenotypically similar (ΔL and θ) shared more outlier nucleotide sites, genes, and biological functions using Mantel tests (Mantel 1967) with 999 permutations.

## Results

### Phenotypic parallelism

We found striking differences between the mean Dune and Headland phenotypes for both plant architecture and leaf characteristics (illustrated in Figure 1C). In multivariate space, Dune and Headland ecotypes clustered into two distinct groups (Figure 2A; Pillai’s Trace = 0.73, F_1, 603_ = 175.13, P < 2.2 × 10^−16^). This pattern held true when traits were separated into plant architecture (Figure S2A; Pillai’s Trace = 0.63, F_1, 603_ = 202.42, P < 2.2 × 10^−16^) and leaf categories (Figure S2B; Pillai’s Trace = 0.61, F_1, 603_ = 233.15, P < 2.2 × 10^−16^). Considering all traits together, the partial effect size of the ecotype term (Wilks partial η^2^ = 0.86) was larger than both the pair (Wilks partial η^2^ = 0.23; Figure S3) and the interaction term (Wilks partial η^2^ = 0.19; Figure 2B). This suggests that the phenotypic variation within the system is mainly explained by differences between ecotypes rather than replicate pairs.

**Figure 2.**
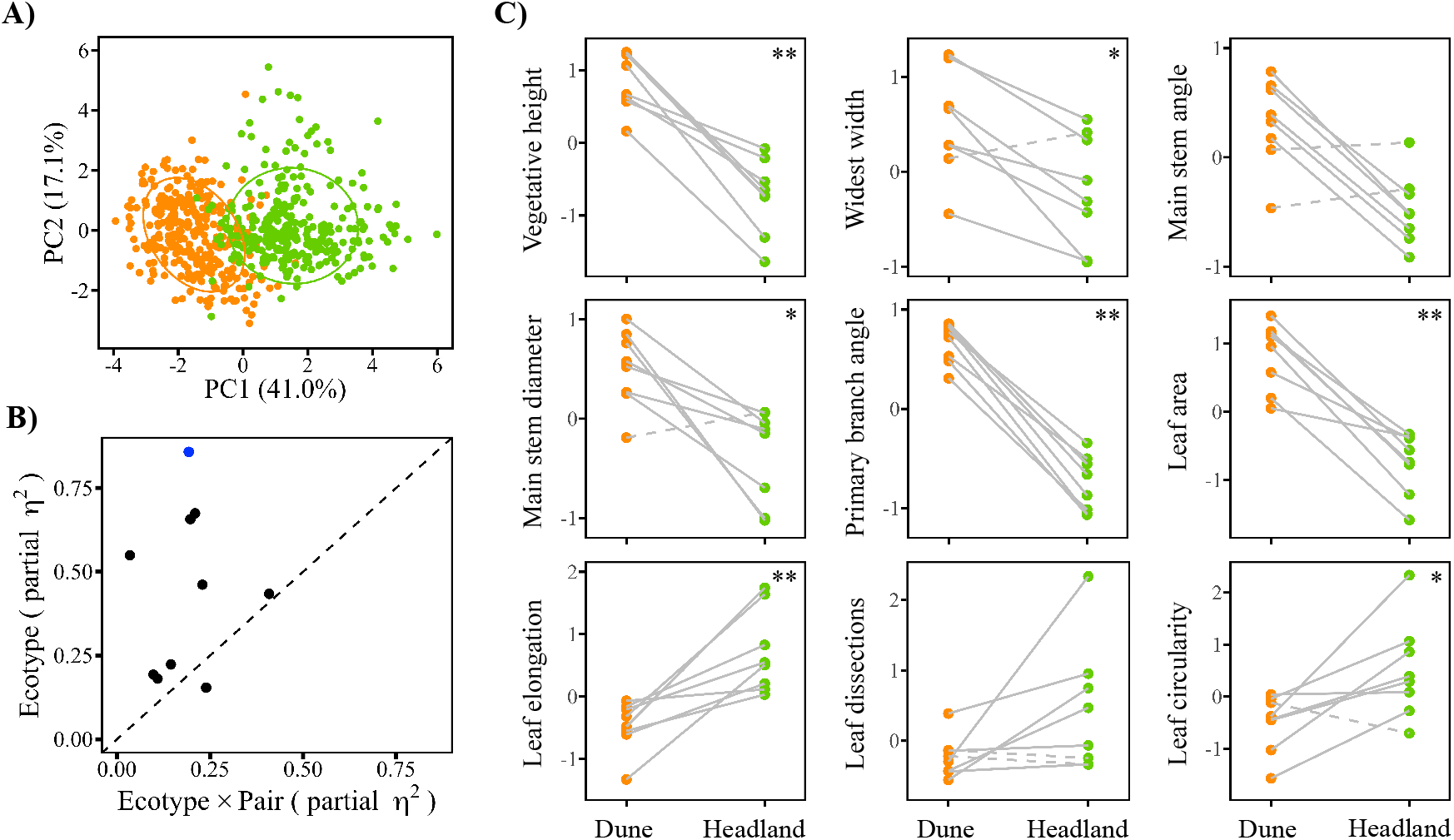
Ecotype and trait phenotypic parallelism. **(A)** Principal component analysis of Dune (orange) and Headland (green) phenotypes (five plant architecture and four leaf traits) across 20 populations. Ecotypes are delimited by 70% probability ellipses. **(B)** Partial effect sizes (partial η^2^) for the ecotype and the interaction (ecotype × pair) for the trait-by-trait linear models, each dot representing a single trait. The blue dot represents Wilk’s partial effect size for all traits combined within the MANOVA. Dashed line is a 1:1 ratio, where points above the line represent a larger contribution of parallel evolution (shared Dune-Headland divergence across localities) than non-parallel evolution (unique Dune-Headland divergence across localities). See Table S5 for exact values. **(C)** Vote-counting for five plant architecture and four leaf traits across eight replicate pairs. Dots represent the mean trait value for each population (N = 30). Lines connect the Dune (orange) populations to their Headland (green) pair at each locality. Dashed lines represent pairs whose Dune-Headland trait value is in the opposite direction from the majority of pairs. Asterisks denote significance (** S-statistic = 8, P = 0.0078; * S-statistic = 7, P = 0.035).

Across all traits, K-means clustering analysis correctly assigned 95% of Dune individuals, and 87% of Headland individuals into the correct cluster, further suggesting most individuals within an ecotype are more phenotypically similar than between ecotypes. When plant architecture and leaf traits were measured separately, these numbers were slightly reduced. For plant architecture traits alone, 91% of Dunes and 82% of Headlands were assigned to the correct cluster. For leaf traits, 93% of Dunes and 78% of Headlands were correctly assigned. We performed a linear discriminant analysis (LDA) on all traits to ask which linear combination of traits best explains the phenotypic differences between Dune and Headland ecotypes. The LDA was strongly loaded by leaf area and primary branch angle, followed by leaf dissection, leaf circularity, and widest width of the plant (Table S5). All traits were loaded in the same direction, except for widest width of the plant, leaf dissection, and leaf circularity. The LDA suggests that divergence between ecotypes is multivariate and has occurred on most measured traits, and that a single trait does not dominate the phenotypic differences between ecotypes.

We also explored the phenotypic differences between the Dune and Headland ecotypes at a univariate trait level. We first used vote-counting to quantify whether the traits in the Dune and Headland populations of each pair have evolved in the same direction. For all traits, at least six of the eight pairs evolved in parallel (Figure 2C). Four of the nine traits had all eight pairs evolving in the same direction (i.e., there was a consistent increase or decrease in the Dune vs Headland mean trait value across replicate pairs; S-statistic = 8, P = 0.0078), and three traits had seven pairs evolving in the same direction (S-statistic = 7, P = 0.035). Trait-by-trait linear models revealed a significant main effect of ecotype for each trait (Table S6), suggesting there are differences between Dune and Headland populations for all traits.

Extracting the effect size for these linear models, the *ecotype* effect size was larger than both the pair (Figure S3) and interaction term (Figure 2B) for most traits (i.e., more data points above the dotted line than below, see Table S6 for details). As observed at the multivariate level, the larger effect sizes for the ecotype terms suggest that the phenotypic variation within the system is mainly explained by differences between ecotypes rather than replicate pairs.

Next, we investigated whether the phenotypic differences between the Dune and Headland of each replicate population pair were consistent across localities using PCVA. Within multivariate phenotypic space, there were different levels of divergence (ΔL) between replicate pairs (Figures 3A and 3B). Considering all traits, the mean ΔL (±SE) between pairs was 1.7 ± 0.15, and out of the 28 pairwise comparisons, we only observed nine statistically parallel comparisons (i.e., ΔL ≈ 0; 32.1% of pairwise comparisons; Table S7). Therefore, most population pairs have different amounts of divergence between the Dune and Headland populations. When we separately analyzed traits as two categories (plant architecture and leaf shape), we captured a signal of parallel divergence across a greater number of replicate pairs (Figures S4 and S5). We observed ten statistically parallel comparisons for plant architecture traits (35.7% of pairwise comparisons; mean ΔL 1.0 ± 0.12; Table S8), and thirteen statistically parallel comparisons for leaf traits (46.4% of pairwise comparisons; mean ΔL 0.93 ± 0.14; Table S9).

**Figure 3.**
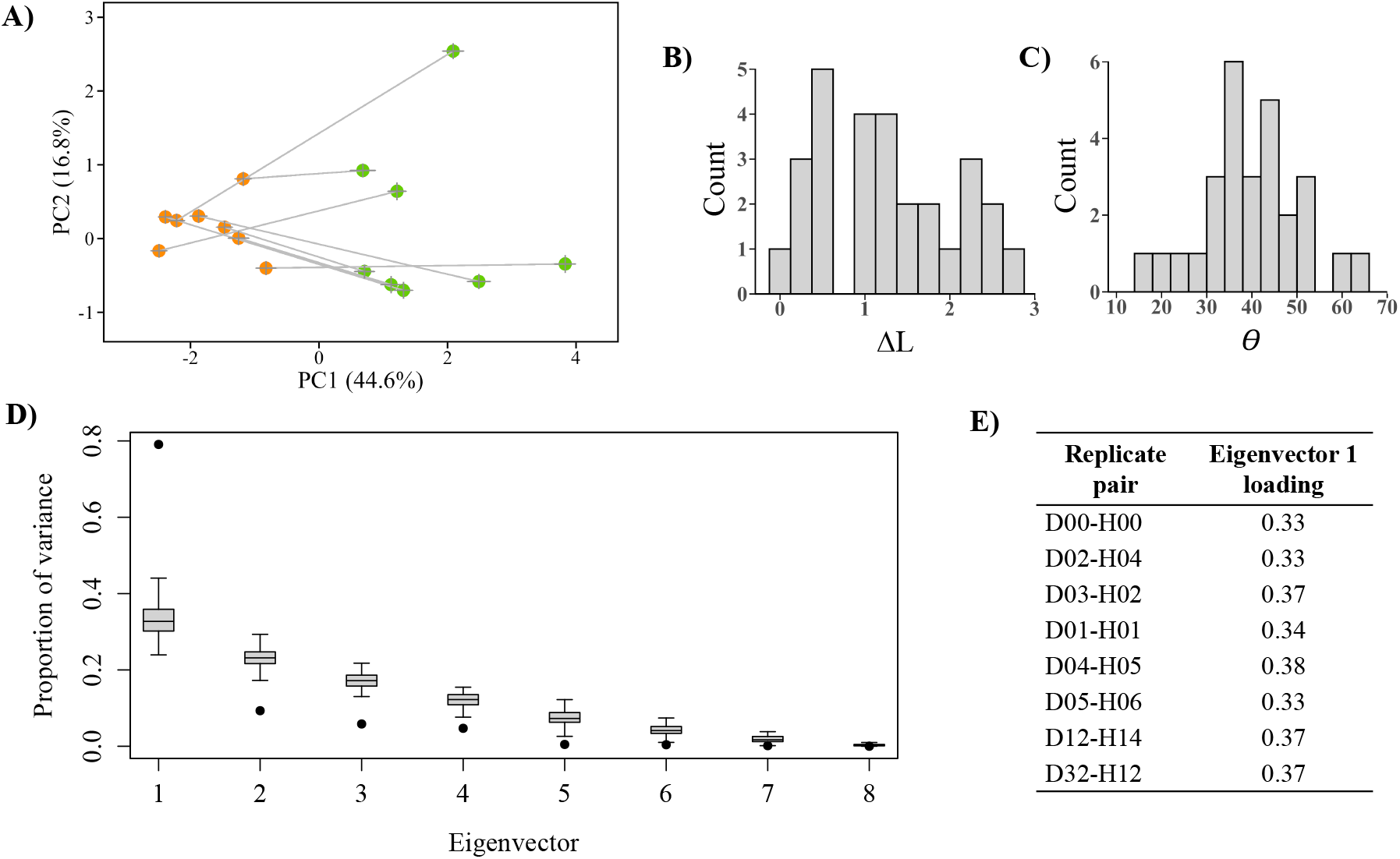
Replicate pair phenotypic parallelism. **(A)** Phenotypic change vector analysis for five plant architecture and four leaf traits across eight replicate Dune-Headland pairs. Each dot represents the population centroid (multivariate phenotypic mean), ± SE. The Dune (orange) and Headland (green) populations of a replicate pair are connected with a line. **(B)** Frequency distribution of the 28 pairwise phenotypic divergences (ΔL) between Dune-Headland replicate pairs (Table S6). **(C)** Frequency distribution of the 28 pairwise contribution of traits (θ) between Dune-Headland replicate pairs (Table S9). **(D)** Proportion of variance across the eight eigenvectors from eigenanalysis of the correlation matrix of the individual pairwise angles between Dune-Headland replicate populations at each locality (Table S7). Grey boxplots represent the null distribution of no shared axes of evolutionary change. **(E)** Loadings of each replicate pair onto the first eigenvector of **(D)**.

The contribution of traits to divergence (θ) was quite variable across pairs (Figures 3A and 3C). Out of the 28 pairwise comparisons, only one angle was parallel, i.e., θ ≈ 0° (3.6% of pairwise comparisons; Table S10), indicating that traits weigh differently in the Dune-Headland divergence across localities. The mean angle (±SE) between population pairs was 39.5 ± 2.1°; all angles were acute, with a maximum of 62.8°. When traits were split into plant architecture and leaf categories, we again captured a stronger signal of phenotypic parallelism for both categories. We observed nine statistically parallel angles for plant architecture traits (mean angle 29.8 ± 3.0°; Table S11) and four statistically parallel angles for leaf traits (42.6 ± 3.4°; Table S12).

We then asked whether there was a shared axis of evolutionary change across replicate pairs by undertaking eigenanalysis on the pairwise angles across localities. We observed that the first eigenvector was the only significant axis, explaining 79% of the phenotypic divergence between ecotypes across localities (Figure 3D). All replicate pairs loaded positively and with similar magnitudes on this first eigenvector, revealing that each replicate pair has evolved in the same direction in multivariate trait space (Figure 3E).

### Genotypic parallelism

#### Parallel nucleotide polymorphisms

Very few sampled SNPs explained more variance between ecotypes than between replicate pairs (Figures 4A, 4C, S6A and S6C). Specifically, only 6.3% of sampled SNPs (607 out of 9,687 SNPs) contained a partial effect size of the ecotype term that was larger than the interaction term (i.e., those above the dashed line in Figure 4A which are also > 0 in Figure 4C). This suggests that, within the dataset presented here, parallel evolution at the level of the nucleotide site within the system is largely predominated by differences between replicate pairs.

**Figure 4.**
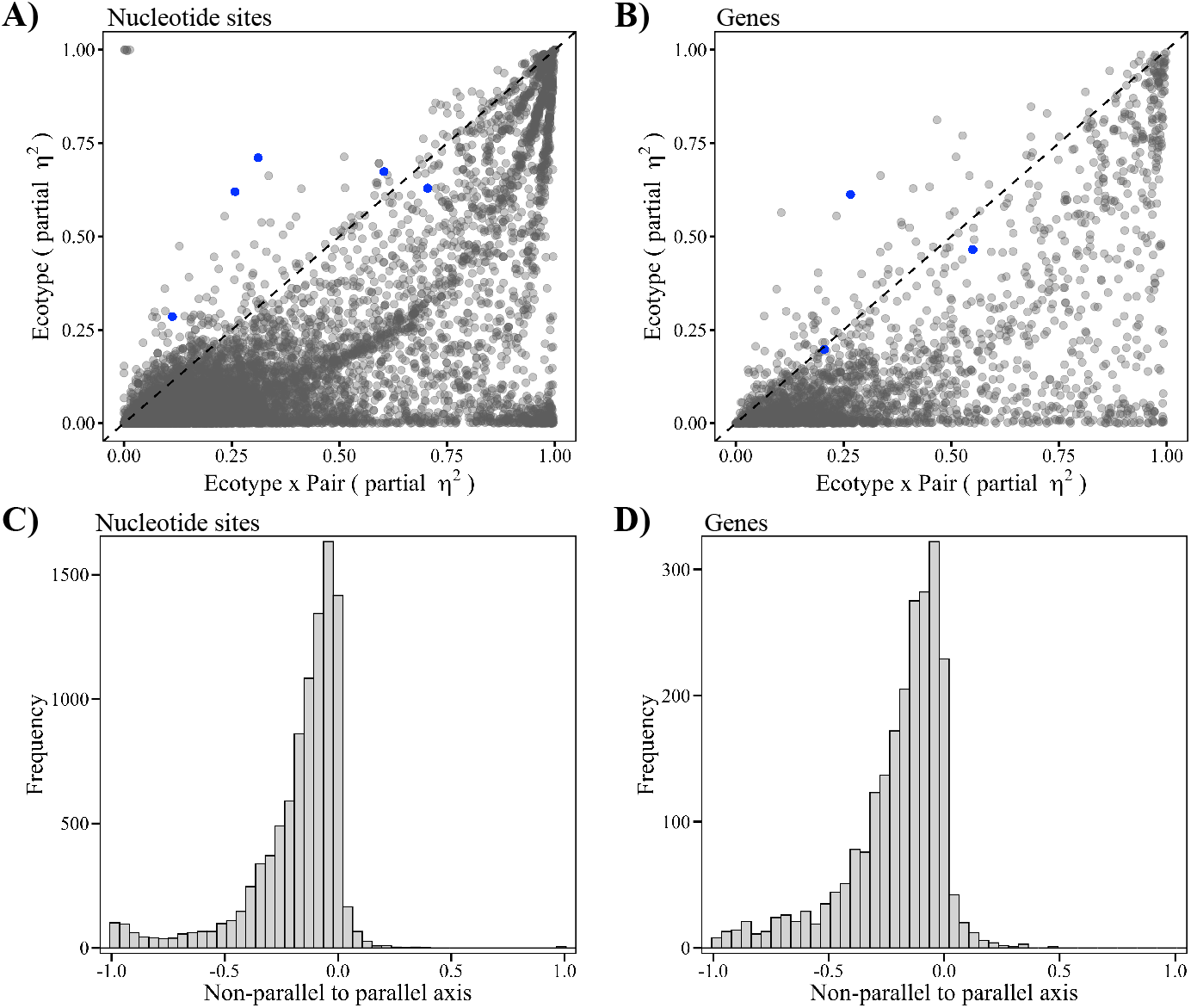
Relative contributions of genotypic parallel and non-parallel evolution. Partial effect sizes (partial η^2^) for the ecotype and the interaction (ecotype *×* pair) from linear models for all sequenced nucleotide sites **(A)** and genes **(B)**. Each dot represents either a single SNP **(A)** or gene **(B)**. Most points fall below the dashed 1:1 ratio line, indicating that the variation in Dune-Headland divergence is largely unique to replicate pairs (non-parallel), rather than shared across localities (parallel). The blue dots denote the best candidates for parallel evolution (those in Figure 5A) at the level of the SNP **(A)** and gene **(B)**. The data in **(A)** and **(B)** is plotted as frequency distributions for the nucleotide sites **(C)** and genes **(D)**. Values represent the distance of the SNP or gene from the 1:1 dashed line of equal effect. Positive values indicate more parallel evolution, whereas negative values indicate more non-parallel evolution. As most values fall below zero, between-ecotype variation at the level of the nucleotide site and gene is mainly unique to replicate pairs.

We identified 93 highly differentiated sites between all Dune and all Headland populations (*Approach 1*; ~1% of sequenced SNPs), with 54 SNPs falling within genic regions and 39 in non-genic regions. For outliers detected separately for the Dune-Headland pairs at each locality (*Approach 2*), there were no outlier SNPs common to all nine pairs. The highest number of pairs with common outlier SNPs was seven pairs, where we detected six SNPs that were outliers (Figure 5B). On average, 157 outlier SNPs (SD = 74.5) were shared between any two localities, and for each of these pairwise comparisons, the shared SNPs were greater than expected by chance (Table S13). We detected 15 nucleotide sites (0.16% of sequenced SNPs; Figure S7) that contained concordant allele frequency differences across localities (*Approach 3*) in either all nine (S-statistic = 9, P = 0.004), or eight (S-statistic = 8, P = 0.04) replicate pairs. Nine of these SNPs fall within genic regions, whereas six are in non-genic regions.

**Figure 5.**
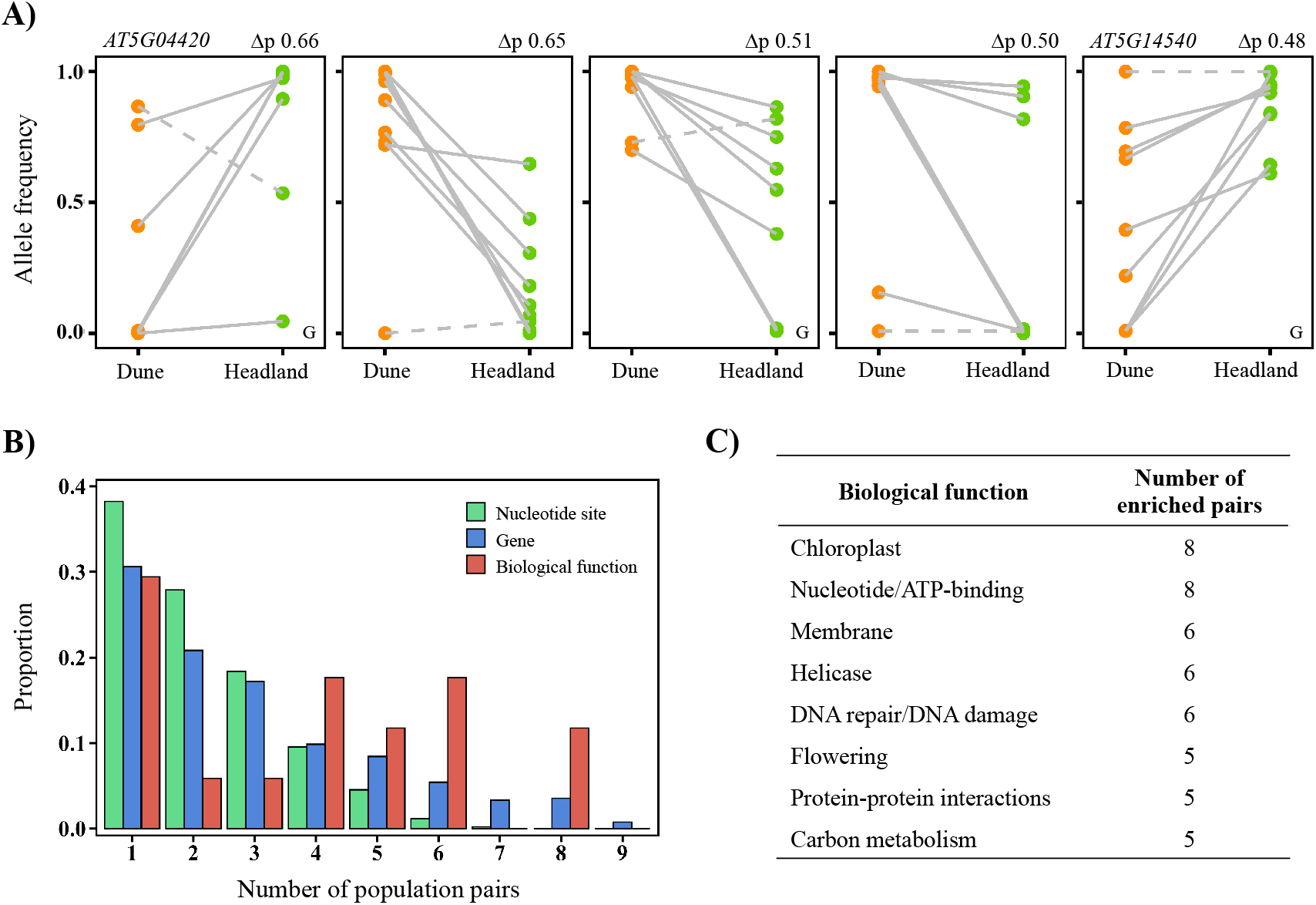
Genotypic parallelism: nucleotide site, gene and biological function. **(A)** Candidate outlier nucleotide sites showing high differentiation between the Dune-Headland ecotypes as well as concordant allele frequency changes across replicate pairs. Dots represent the allele frequency value (of the reference allele) for each population. Lines connect the Dune (orange) populations to their Headland (green) pair at each locality. Dashed lines represent pairs whose Dune-Headland change in allele frequency is in the opposite direction from the majority of pairs. Δ*p* denotes the overall change in allele frequency between the ecotypes. *G* denotes SNPs that occur within genic regions. **(B)** Proportion of outlier nucleotide sites, outlier genes, and enriched biological functions shared across the nine replicate pairs. **(C)** Enriched biological functions shared across five or more replicate population pairs.

As we did not detect any outliers across all nine or eight replicate pairs in *Approach 2*, we consider our best parallel SNP candidates across the *S. lautus* system using only *Approach 1* and *3*. Five SNPs were detected as outliers in *Approach 1* and *3* (three genic, two non-genic; Figure 5A; red dots in Figures 4A), showing high differentiation between ecotypes, with concordant allele frequency changes across localities. The average difference in allele frequency between Dune and Headlands for the three genic SNPs was 0.55 (SD = 0.096), whereas the average for the two non-genic SNPs was 0.57 (SD = 0.107).

#### Parallel genic polymorphisms

Very few genes explained more variance between ecotypes than between pairs (Figures 4B, 4D, S6B and S6D). More specifically, only 5.6% of genes (148 out of 2320 genes) contained a partial effect size of the ecotype term that was larger than the interaction term (i.e., those above the dashed line in Figure 4A which are > 0 in Figure 4C). This indicates that there is more parallel evolution at the level of the gene than the SNP, and that parallelism at the level of the gene is largely predominated by differences between replicate pairs.

Of the five candidate outlier SNPs identified above using the outlier approach (i.e., those showing high differentiation between ecotypes in *Approach 1* and concordant allele frequency chances across replicate pairs in *Approach 3*), the three genic SNPs fall within three separate genes, two of which have homologs within *Arabidopsis* (Table S14; red dots in Figures 4B). These two genes encode a galactose oxidase/kelch repeat superfamily protein (*AT5G04420*; Figure 5A first panel) and a basic salivary proline-rich-like protein (*AT5G14540*; Figure 5A last panel). The proteins are both located in the cytosol and are both expressed in a wide variety of tissue types (Klepikova et al. 2016). Considering *Approach 1* and *3* separately, the 54 outlier genic SNPs detected in *Approach 1* fall in 49 separate genes, of which 44 have homologs within *Arabidopsis*. The majority of these genes are involved in processes including ion transport, transcription, response to heat, response to water deprivation, DNA repair and embryo development (see Table S14 for details of each gene). For *Approach 3*, the nine outlier genic SNPs from fall in nine separate genes, of which seven have homologs within *Arabidopsis*. These genes are involved in processes including ion transport, aminoacylation, embryo development and DNA repair (see Table S14 for details of each gene).

We detected highly differentiated genes between the Dune and Headland of each locality (*Approach 2*), and compared how many of these outlier genes were common between all pairwise comparisons of replicate pairs. On average, 124 outlier genes (SD = 54.6) were shared between any two replicate pairs. The shared outlier genes between all pairwise comparisons were greater than expected by chance, except for one comparison (D14-H15 vs D32-H12; Table S14). Thirty-nine genes were outliers in at least eight or nine replicate pairs (Figure 5B), of which 36 contained homologs in *Arabidopsis*. These genes are involved in processes including ion transport, transcription, seed development, response to auxin, response to heat, response to salt stress, embryo development and cell growth (see Table S14 for details of each gene).

#### Enriched biological functions

For each replicate pair we conducted a gene-enrichment analysis using outlier genes to ask whether any biological functions were enriched. Examining individual replicate pairs, we detected a total of 17 enriched functions (Figure 5C; Table S16). However, no function was repeatedly enriched in all nine replicate pairs, although two functions (*chloroplast* and *nucleotide-binding/ATP-binding;* UniProtKB keywords) were repeatedly enriched across eight replicate pairs (S-statistic = 8, P = 0.04). Within the *chloroplast* category, most outlier genes across pairs and are involved in processes including oxidation reduction, response to light, translation, proteolysis, protein phosphorylation and protein folding. See Table S17 for details on each gene within the *chloroplast* category and the number of replicate pairs the genes were detected as an outlier. Within the *nucleotide-binding*/*ATP-binding* category, most outlier genes across pairs are involved in processes including protein phosphorylation, protein folding, transcription, aminoacylation, ion transport, and response to stress. See Table S18 for details on each gene within the *nucleotide*/*ATP-binding* category and the number of replicate pairs the genes were detected as an outlier.

Finally, we examined the distributions of the shared outlier nucleotide sites, outlier genes, and enriched biological functions across the nine replicate pairs. We observed. (i.e., a distribution more skewed to the right, suggestive of more parallelism at the level of the biological function compared to the SNP or gene.

We further asked whether the distributions of shared outlier nucleotide sites, outlier genes, and enriched biological functions across the nine replicate pairs were significantly different. The distribution of the proportion of shared outlier nucleotide sites was significantly different to the outlier genes (X^2^ = 279.65, df = 8, P < 2.2×10^−16^), and biological functions (X^2^ = 361.95, df = 8, P < 2.2×10^−16^). The distributions of the genes and biological functions were not significantly different across the nine replicate pairs (X^2^=14.52, df=8, P = 0.069).

### Variation in phenotypic parallelism

Gene flow did not constrain phenotypic divergence. There was no relationship between levels of gene flow and the lengths of phenotypic vectors (L) between ecotypes within a locality when considering (i) Dune to Headland gene flow (F_1,5_ = 1.67, P = 0.25, R^2^ = 0.25), Headland to Dune gene flow (F_1,5_ = 3.44, P = 0.12, R^2^ = 0.41), or (iii) average gene flow (F_1,5_ = 2.29, P = 0.19, R^2^ = 0.31). Moreover, we did not find a relationship between divergence time between ecotypes and L within a locality (F_1,5_ = 1.04, P = 0.35, R^2^ = 0.17). Environmental distance did not relate to how phenotypically divergent (L) a population pair was (F_1,3_ = 0.046, P = 0.84, R^2^ = 0.015), although we treat these data with caution as environmental data were only available for five localities. Population pairs that were more phenotypically similar (i.e., smaller ΔL) did not share more outlier SNPs, genes or biological functions than those with large ΔL (Mantel test SNPs: r = −0.215, P = 0.894; Mantel test genes: r = −0.164, P = 0.835, Mantel test biological functions: r = −0.179, P = 0.837). Population pairs with similar contribution of traits to divergence (i.e., smaller θ) also did not share more outlier SNPs, genes, or biological functions (Mantel test SNPs: r = −0.493, P = 0.989; Mantel test genes: r = −0.347, P = 0.865; Mantel test biological functions: r = −0.337, P = 0.93). These results imply that, with the current genetic data, the extent of phenotypic parallelism in the system is not largely driven by the underlying genetics or demographic history.

## Discussion

Understanding the way in which independent populations adapt to their environment when faced with similar selective pressures, allows us to gain insight into the repeatability and predictability of evolution. Here, we have demonstrated striking phenotypic parallel evolution in the highly replicated *Senecio lautus* system. Multiple instances of adaptation to parapatric Dune and Headland environments have consistently resulted in the repeated evolution of ecotypes with contrasting morphologies. Although there is some variation between ecotypic divergences across localities, all replicate pairs follow a common evolutionary trajectory within phenotypic space. Across replicate localities, Dune and Headland ecotypes have diverged mainly via mutational changes in different genes, although some of these belong to the same predicted biological function. This implies that evolution within the *S. lautus* system may be somewhat flexible at lower levels of biological organization, yet more constrained at the functional level. Given the high levels of evolutionary independence amongst populations with similar phenotypes (James et al. 2021), our current work positions *S. lautus* as an ideal candidate to examine how adaptive replicated evolution arises in nature.

### Phenotypic parallelism

Phenotypic variance in the *S. lautus* system is explained mostly by the differences between ecotypes rather than the replicate pairs, indicating that the phenotypic differences between ecotypes are consistent across localities. There is also clear phenotypic separation between Dune and Headland *S. lautus* ecotypes in the first two dimensions of multivariate trait space. This distinct separation of ecotypes is seen in other empirical parallel evolution systems such as lake-stream stickleback on Haida Gwaii in Canada (Deagle et al. 2012), and dwarf-normal lake whitefish (Laporte et al. 2015). In contrast, systems such as benthivorous-planktivorous Arctic charr (Jacobs et al. 2020), benthic-limnetic cichlid fishes (Elmer et al. 2014) and lake-stream threespine stickleback on Vancouver Island in Canada (Stuart et al. 2017) have large overlap between ecotypes in phenotypic space, revealing that the between-ecotypic divergences are variable across replicate pairs, i.e., that evolution is to some degree of non-parallel. In contrast, the phenotypically distinct Dune and Headland *S. lautus* ecotypes position the system as highly parallel and repeatable at the multivariate trait level.

At the univariate level, most traits show consistent differences between Dune and Headland ecotypes across replicate localities. The trait that displayed the greatest non-parallelism was leaf dissection, which varied more between replicate pairs than between ecotypes, though this trait still showed consistent trait differences between ecotypes in six out of eight pairs. Thus, even though there is some Dune-Headland trait variation between localities, all *S. lautus* traits are highly parallel. This pattern is rather different to other systems such as threespine stickleback. Here, Stuart et al. (2017) found that most trait variance is explained by differences between replicate pairs rather than between ecotypes, suggesting little parallelism at the level of individual traits. Although this pattern might occur in *S. lautus* if more traits are measured, the phenotypic dimensionality of the system does not seem to be very high: of the 14 traits we measured, we discarded five highly correlated traits. In other studies of *S. lautus* (Walter et al. 2018a), strong genetic correlations exist amongst a variety of vegetative traits suggesting strong interdependence between morphological modules such as leaf and plant architecture, and high genetic constraint in the system. In the current work although we lack the ability to make inferences of the exact traits under the direct target of selection, we have likely measured these and correlated traits which together reveal the overall phenotypic differences of populations and ecotypes in the system. It will be interesting for future work to further explore the effect of correlated selection on our ability to measure replicated evolution and its contribution to differences amongst replicates.

Pairwise comparisons of Dune-Headland phenotypic divergence (PCVA) revealed different magnitudes of divergence and different contribution of traits to divergence for most pairs. Yet, surprisingly, phenotypic divergence shares a common multivariate evolutionary trajectory in the system: there was only one significant axis of evolutionary change that explained 78% of the phenotypic variance across the system. All *S. lautus* replicate pairs have the same shared evolutionary trajectory in this single dimension of multivariate space (i.e., they all load with the same direction and similar magnitude on this first eigenvector). This is in contrast to the classic lake-stream stickleback system, where multiple dimensions of evolutionary change are significant, and replicate lineages have not all evolved along the same trajectory (i.e., load with opposing signs onto the significant eigenvectors; De Lisle and Bolnick 2020). Phenotypic parallel evolution in *S. lautus* is therefore considered ‘complete’, as the Dune-Headland phenotypic divergence at each locality have evolved in the same way (De Lisle and Bolnick 2020).

### Genotypic parallelism

Within *S. lautus*, genotypic parallelism was strongest at the level of the biological function. Although there was some shared SNP and gene outliers between pairwise localities, we only detected five candidate outlier nucleotide sites that were parallel across the entire system. These results suggest that adaptation in *S. lautus* is flexible at lower levels of organization (the nucleotide site and gene), and more constrained at the level of the biological function. Non-parallelism at the level of the SNP and gene suggests there is a large amount of genetic redundancy in the *S. lautus* system (Barghi et al. 2020), where different genetic routes can produce adaptive phenotypes with similar fitness (Láruson et al. 2020). This non-parallelism is also a characteristic signal of polygenic adaptation (Barghi et al. 2020), where adaptation proceeds via many alleles of small effect (Yeaman 2015). In such systems where adaptation occurs via different genetic routes in replicate populations, it is perhaps not surprising that we detect little parallelism at the SNP and gene. Furthermore, parallelism at the biological function level might be common in both plants and animals (e.g., Smith and Rausher 2011; Kowalko et al. 2013; Roda et al. 2013b; Laporte et al. 2015; Perreault-Payette et al. 2017; Cassin-Sackett et al. 2019), given that there are fewer biological functions than there are genes or nucleotide sites (Tenaillon et al. 2012; Tiffin and Ross-Ibarra 2014).

Recent studies in *S. lautus* have demonstrated that hormone signaling, specifically the auxin pathway, is divergent between Dune and Headland populations (Roda et al. 2013b; Wilkinson et al. 2021). Auxin plays a key role in a plants ability to respond to gravity (Strohm et al. 2012), and is strongly correlated with the prostrate and erect growth forms within the system (Wilkinson et al. 2021). We therefore expected to find highly differentiated auxin-related genes within our current study. Consistent with this prediction, we detected divergent genes involved in the auxin pathway that are differentiated across multiple population pairs, including *GH3.1* (Staswick et al. 2005), *NPH4* (Harper et al. 2000) and genes from the ABCB family (Cho and Cho 2013; see Table S19 and Supplementary Methods S4 for more details). This gives further evidence that auxin may play a key role in creating the contrasting growth habits in *S. lautus*. Future studies on the molecular basis of adaptation should focus on the concomitant contribution of many genes to phenotypic variation and to their shared cellular and physiological roles, as it is likely that variation in regulatory networks might underlie a large fraction of the adaptive space in organisms (Boyle et al. 2017; VanWallendael et al. 2019).

### The nature of parallel evolution in S. lautus

Empirical systems of parallel evolution allow us to address how repeatable and predictable evolution is within nature, yet many factors can cause deviations between replicate populations. For instance, demographic history, environmental heterogeneity within each habitat, the interplay between the genotype, phenotype and fitness landscapes, genetic constraints, and stochastic forces such as genetic drift can all impact the likelihood of parallel evolution across replicate localities (Lenormand et al. 2009, 2016; Conte et al. 2012; Rosenblum et al. 2014; Ord and Summers 2015; Fraïsse and Welch 2019). The clear trait-environment association observed within the *S. lautus* system is quite remarkable, despite varying levels of gene flow, divergence times (James et al. 2021), environmental distances (Roda et al. 2013b) and selection largely acting upon different SNPs and genes between parapatric ecotypes across localities. As every instance of repeated Dune-Headland evolution has resulted in extremely similar phenotypic adaptations, we can the adopt the simple, binary classification of ‘Dune’ and ‘Headland’ to describe the ecotypes (De Lisle and Bolnick 2020). This is surprisingly not the case for many systems of parallel evolution including one of the most famous cases in nature, the threespine stickleback (De Lisle and Bolnick 2020). Replicate lake-stream stickleback populations show a large degree of non-parallel evolution, implying that categorical terms of ‘lake’ and ‘stream’ might misrepresent the large degree of phenotypic overlap between ecotypes.

Although our current work has revealed that the Dunes and Headlands are quite phenotypically distinct, there are still phenotypic differences between the populations within each ecotype. This seems to be more pronounced in the Headlands (the phenotypic centroids of Dune populations cluster, but Headland populations are somewhat more scattered in multivariate space). This might be explained by the nature of correlated selection in each ecotype which can lead to different phenotypic and fitness landscapes in *S. lautus*. Previous reciprocal transplant experiments have demonstrated that ecotypes are locally adapted and exhibit a strong reduction in fitness when grown in foreign habitats (Melo et al. 2014; Richards and Ortiz-Barrientos 2016; Richards et al. 2016; Walter et al. 2016, 2018b; Wilkinson et al. 2021). However, different to Dune individuals that are equally fit across other non-local sand dune habitats, Headland individuals have reduced fitness in non-local headland habitats (Walter et al. 2016). These observations suggest some environmental heterogeneity within rocky headlands that can affect correlated selection in each locality. The fitness landscape for Headlands might therefore be either broad and rugged, or with multiple optima, with each Headland population residing on a different local optimum. Overall, these differences in fitness landscapes within each ecotype may reflect why we see some phenotypic variation between Dune-Headland pairs across localities, yet stabilised within specific multivariate trajectories.

### The effects of sampling on parallelism

The ability to detect genotypic parallelism is impacted by sampling. Reduced representation libraries, such as those used within the current work, sparsely sample the genome and will likely fail to detect many loci involved in adaptation (Tiffin and Ross-Ibarra 2014; Lowry et al. 2017). This is of particular concern when adaptation occurs via few genes of large effect (e.g., the *Eda* gene in sticklebacks; Colosimo et al. 2005), as these genes will likely not be sampled in the absence of whole genome sequencing. In contrast, adaptation within *S. lautus* seems to be polygenic (Yeaman 2015), being underpinned by the frequency shift of many different alleles and genes across replicate localities (also see Roda et al. 2013b; Wilkinson et al. 2021). Our reduced representation libraries will still likely capture a proportion of the many variants involved in adaptation, although we acknowledge that our current work focuses on highly divergent alleles and we have disregarded alleles with subtle changes in allele frequencies. Rather than placing an emphasis on the specific genes involved in parallel evolution, we have demonstrated that genotypic evolution is likely more parallel at higher levels of biological organization. Future whole genome sequencing will help elucidate the relative contributions of all variants to adaptive evolution, including those with small effects (Barghi et al. 2020).

Local variation in genetic divergence is impacted by genome features including recombination rates (Booker et al. 2020), background selection, linkage and demography (see Hoban et al. (2016) for a review). This impacts our ability to detect regions involved in replicated divergence. For instance, linked selection in regions of low recombination could increase divergence relative to neutral expectations in all populations examined and not only in a specific parapatric pair. In other words, evolutionary constraints might lead to signals of replicated evolution, when in fact they inexorably arise as consequence selection interacting with conserved genomic features and unrelated to speciation events. Additionally, within this study we have not examined other aspects of the genome that can be involved in adaptation including copy number variation (Schrider et al. 2016; Nelson et al. 2019), inversions (Kirkpatrick and Barton 2006; Lowry and Willis 2010; Faria et al. 2019), transposons (González and Petrov 2009; Schrader and Schmitz 2019), and variation in gene expression levels (Rivas et al. 2018; Verta and Jones 2019). Future work examining these aspects of parallel evolution will help us gain a more complete picture of the dynamics of phenotypic parallelism within *S. lautus*. Nevertheless, our current genetic dataset reveals that highly parallel phenotypes need not arise due to the exact same underlying genetic mechanisms, which supports the findings of other reduced representation datasets in the *S. lautus* system (Roda et al. 2013b; Wilkinson et al. 2021).

Finally, we must be aware that researchers of replicated evolution often implement different statistical approaches to measure parallelism across systems. These different approaches can lead to different interpretations of parallel evolution both at the level of the phenotype and genotype (Bolnick et al. 2018). It is thus evident that the field requires progress towards a common framework to allow researchers to quantify and compare the exact extent of parallelism across systems, considering that parallelism can manifest at different scales as well as different levels of biological organization. For instance, further work needs to enrich current theories of multi-trait evolution so we can develop better null hypotheses for parallel evolution while accounting for correlations between traits, including those that are highly pleiotropic (Yeaman 2015; De Lisle and Bolnick 2020). Furthermore, to have a more complete understanding of the dynamics and link between genotypic and phenotypic parallel evolution, studies should aim to identify causal mutations across replicate populations and to ask whether any shared variants have arisen via de novo mutations, standing genetic variation or adaptive introgression. It is necessary to then directly link variants to adaptive traits and further demonstrate the traits confer a fitness advantage to native populations.

### Conclusions

Overall, we have demonstrated that the highly replicated Dune-Headland *S. lautus* system is a remarkable case of parallel phenotypic evolution in nature. Independent populations have repeatedly evolved extremely similar phenotypes during the adaptation to coastal environments. Genotypic divergence has largely occurred via different mutations in different genes across replicate populations, implying that evolution in the system is highly polygenic. The enrichment of similar biological functions across replicate localities suggests that genotypic adaptation may be constrained at higher levels of biological organization. The *S. lautus* system allows us to examine the repeatability and predictability of evolution, and understand how genetic redundancy and functional constraint impact the likelihood of parallel evolution within natural systems.

## Acknowledgements

We are grateful to HN Nguyen Vu for assisting with sample collection. We thank SI De Lisle and Ortiz-Barrientos laboratory members for insightful comments on previous versions of this manuscript. S Chenoweth and M Blows provided very useful feedback on ME James’ PhD dissertation. This work was supported by an Australian Research Council grant (DP190103039) to DO and JE, and a University of Queensland Graduate School International Travel Award to MEJ.

## Notes

### Competing Interest Statement

The authors have declared no competing interest.

## References

Adams, D. C., and M. L. Collyer. 2009. A general framework for the analysis of phenotypic trajectories in evolutionary studies. Evolution 63:1143–1154.

Altschul, S. F., W. Gish, W. Miller, E. W. Myers, and D. J. Lipman. 1990. Basic local alignment search tool. J. Mol. Biol. 215:403–410.

Arendt, J., and D. Reznick. 2008. Convergence and parallelism reconsidered: what have we learned about the genetics of adaptation? Trends Ecol. Evol. 23:26–32.

Barghi, N., J. Hermisson, and C. Schlötterer. 2020. Polygenic adaptation: a unifying framework to understand positive selection. Nat. Rev. Genet., doi: 10.1038/s41576-020-0250-z.

Blount, Z. D., R. E. Lenski, and J. B. Losos. 2018. Contingency and determinism in evolution: Replaying life’s tape. Science 362:eaam5979.

Bohutínská, M., J. Vlček, S. Yair, B. Laenen, V. Konečná, M. Fracassetti, T. Slotte, and F. Kolář. 2021. Genomic basis of parallel adaptation varies with divergence in *Arabidopsis* and its relatives. PNAS 118:e2022713118.

Bolnick, D. I., R. D. H. Barrett, K. B. Oke, D. J. Rennison, and Y. E. Stuart. 2018. (Non)Parallel evolution. Annu. Rev. Ecol. Evol. Syst. 49:303–330.

Booker, T. R., S. Yeaman, and M. C. Whitlock. 2020. Variation in recombination rate affects detection of outliers in genome scans under neutrality. Mol. Ecol. 29:4274–4279.

Boyle, E. A., Y. I. Li, and J. K. Pritchard. 2017. An expanded view of complex traits: From polygenic to omnigenic. Cell 169:1177–1186.

Butlin, R. K., M. Saura, G. Charrier, B. Jackson, C. André, A. Caballero, J. A. Coyne, J. Galindo, J. W. Grahame, J. Hollander, P. Kemppainen, M. Martínez-Fernández, M. Panova, H. Quesada, K. Johannesson, and E. Rolán-Alvarez. 2014. Parallel evolution of local adaptation and reproductive isolation in the face of gene flow. Evolution 68:935–949.

Cai, Z., L. Zhou, N. N. Ren, X. Xu, R. Liu, L. Huang, X.-M. Zheng, Q.-L. Meng, Y.-S. Du, M.-X. Wang, M.-F. Geng, W.-L. Chen, C.-Y. Jing, X.-H. Zou, J. Guo, C.-B. Chen, H.-Z. Zeng, Y.-T. Liang, X.-H. Wei, Y.-L. Guo, H.-F. Zhou, F.-M. Zhang, and S. Ge. 2019. Parallel speciation of wild rice associated with habitat shifts. Mol. Biol. Evol. 36:875–889.

Cassin-Sackett, L., T. E. Callicrate, and R. C. Fleischer. 2019. Parallel evolution of gene classes, but not genes: Evidence from Hawai’ian honeycreeper populations exposed to avian malaria. Mol. Ecol. 28:568–583.

Chevin, L. M., G. Martin, and T. Lenormand. 2010. Fisher’s model and the genomics of adaptation: restricted pleiotropy, heterogenous mutation, and parallel evolution: heterogeneous mutation effects across loci. Evolution 64:3213–3231.

Cho, M., and H. Cho. 2013. The function of ABCB transporters in auxin transport. Plant Signal. Behav. 8:e22990.

Christin, P. A., D. M. Weinreich, and G. Besnard. 2010. Causes and evolutionary significance of genetic convergence. Trends Genet. 26:400–405.

Collyer, M. L., and D. C. Adams. 2007. Analysis of two-state multivariate phenotypic change in ecological studies. Ecology 88:683–692.

Collyer, M. L., D. J. Sekora, and D. C. Adams. 2015. A method for analysis of phenotypic change for phenotypes described by high-dimensional data. Heredity 115:357–365.

Colosimo, P. F., K. E. Hosemann, S. Balabhadra, G. Villarreal, M. Dickson, J. Grimwood, J. Schmutz, R. M. Myers, D. Schluter, and D. M. Kingsley. 2005. Widespread parallel evolution in sticklebacks by repeated fixation of Ectodysplasin alleles. Science 307:1928–1933.

Conte, G. L., M. E. Arnegard, C. L. Peichel, and D. Schluter. 2012. The probability of genetic parallelism and convergence in natural populations. Proc. R. Soc. B 279:5039–5047.

De Lisle, S. P., and D. I. Bolnick. 2020. A multivariate view of parallel evolution. Evolution 74:1466–1481.

Deagle, B. E., F. C. Jones, Y. F. Chan, D. M. Absher, D. M. Kingsley, and T. E. Reimchen. 2012. Population genomics of parallel phenotypic evolution in stickleback across stream–lake ecological transitions. Proc. R. Soc. B 279:1277–1286.

Elmer, K. R., S. Fan, H. Kusche, M. Luise Spreitzer, A. F. Kautt, P. Franchini, and A. Meyer. 2014. Parallel evolution of Nicaraguan crater lake cichlid fishes via non-parallel routes. Nat. Commun. 5:5168.

Elmer, K. R., H. Kusche, T. K. Lehtonen, and A. Meyer. 2010. Local variation and parallel evolution: morphological and genetic diversity across a species complex of neotropical crater lake cichlid fishes. Philos. Trans. R. Soc. B 365:1763–1782.

Elmer, K. R., and A. Meyer. 2011. Adaptation in the age of ecological genomics: insights from parallelism and convergence. Trends Ecol. Evol. 26:298–306.

Faria, R., K. Johannesson, R. K. Butlin, and A. M. Westram. 2019. Evolving inversions. Trends Ecol. Evol. 34:239–248.

Foll, M., and O. Gaggiotti. 2008. A genome-scan method to identify selected loci appropriate for both dominant and codominant markers: A Bayesian perspective. Genetics 180:977–993.

Foster, S. A., G. E. McKinnon, D. A. Steane, B. M. Potts, and R. E. Vaillancourt. 2007. Parallel evolution of dwarf ecotypes in the forest tree *Eucalyptus globulus*. New Phytol. 175:370–380.

Fox, J., M. Friendly, and G. Monette. 2018. heplots: Visualizing Tests in Multivariate Linear Models. R package version 1.3-5. https://CRAN.R-project.org/package=heplots.

Fraïsse, C., and J. J. Welch. 2019. The distribution of epistasis on simple fitness landscapes. Biol. Lett. 15:20180881.

González, J., and D. A. Petrov. 2009. The adaptive role of transposable elements in the Drosophila genome. Gene 448:124–133.

Haas, O., and G. G. Simpson. 1946. Analysis of some phylogenetic terms, with attempts at redefinition. Proc. Am. Philos. Soc. 90:319–349.

Harper, R. M., E. L. Stowe-Evans, D. R. Luesse, H. Muto, K. Tatematsu, M. K. Watahiki, K. Yamamoto, and E. Liscum. 2000. The *NPH4* locus encodes the auxin response factor ARF7, a conditional regulator of differential growth in aerial *Arabidopsis* tissue. Plant Cell 12:757–770.

Hartigan, J. A., and M. A. Wong. 1979. A K-Means clustering algorithm. Appl. Stat. 28:100.

Hoban, S., J. L. Kelley, K. E. Lotterhos, M. F. Antolin, G. Bradburd, D. B. Lowry, M. L. Poss, L. K. Reed, A. Storfer, and M. C. Whitlock. 2016. Finding the genomic basis of local adaptation: pitfalls, practical solutions, and future directions. Am. Nat. 188:379–397.

Huang, D. W., B. T. Sherman, and R. A. Lempicki. 2009a. Bioinformatics enrichment tools: paths toward the comprehensive functional analysis of large gene lists. Nucleic Acids Res. 37:1–13.

Huang, D. W., B. T. Sherman, and R. A. Lempicki. 2009b. Systematic and integrative analysis of large gene lists using DAVID bioinformatics resources. Nat. Protoc. 4:44–57.

Jacobs, A., M. Carruthers, A. Yurchenko, N. V. Gordeeva, S. S. Alekseyev, O. Hooker, J. S. Leong, D. R. Minkley, E. B. Rondeau, B. F. Koop, C. E. Adams, and K. R. Elmer. 2020. Parallelism in eco-morphology and gene expression despite variable evolutionary and genomic backgrounds in a Holarctic fish. PLoS Genet. 16:e1008658.

James, M. E., H. Arenas-Castro, J. S. Groh, S. L. Allen, J. Engelstädter, and D. Ortiz-Barrientos. 2021. Highly replicated evolution of parapatric ecotypes. Accepted Mol. Biol. Evol., doi: 10.1101/2020.02.05.936401.

Jones, F. C., Y. F. Chan, P. Russell, E. Mauceli, J. Johnson, R. Swofford, M. Pirun, M. C. Zody, S. White, E. Birney, S. Searle, J. Schmutz, J. Grimwood, M. C. Dickson, R. M. Myers, C. T. Miller, B. R. Summers, A. K. Knecht, S. D. Brady, H. Zhang, A. A. Pollen, T. Howes, C. Amemiya, E. S. Lander, F. Di Palma, K. Lindblad-Toh, and D. M. Kingsley. 2012. The genomic basis of adaptive evolution in threespine sticklebacks. Nature 484:55–61.

Kirkpatrick, M., and N. Barton. 2006. Chromosome inversions, local adaptation and speciation. Genetics 173:419–434.

Klepikova, A. V., A. S. Kasianov, E. S. Gerasimov, M. D. Logacheva, and A. A. Penin. 2016. A high resolution map of the *Arabidopsis thaliana* developmental transcriptome based on RNA-seq profiling. Plant J. 88:1058–1070.

Knotek, A., V. Konečná, G. Wos, D. Požárová, G. Šrámková, M. Bohutínská, V. Zeisek, K. Marhold, and F. Kolář. 2020. Parallel alpine differentiation in *Arabidopsis arenosa*. Front. Plant Sci. 11:561526.

Konečná, V., M. D. Nowak, and F. Kolář. 2019. Parallel colonization of subalpine habitats in the central European mountains by *Primula elatior*. Sci. Rep. 9:3294.

Kowalko, J. E., N. Rohner, T. A. Linden, S. B. Rompani, W. C. Warren, R. Borowsky, C. J. Tabin, W. R. Jeffery, and M. Yoshizawa. 2013. Convergence in feeding posture occurs through different genetic loci in independently evolved cave populations of *Astyanax mexicanus*. PNAS 110:16933–16938.

Langerhans, R. B., and T. J. DeWitt. 2004. Shared and unique features of evolutionary diversification. Am. Nat. 164:335–349.

Laporte, M., S. M. Rogers, A.-M. Dion-Côté, E. Normandeau, P.-A. Gagnaire, A. C. Dalziel, J. Chebib, and L. Bernatchez. 2015. RAD-QTL mapping reveals both genome-level parallelism and different genetic architecture underlying the evolution of body shape in lake whitefish (*Coregonus clupeaformis*) species pairs. G3 5:1481–1491.

Láruson, Á. J., S. Yeaman, and K. E. Lotterhos. 2020. The importance of genetic redundancy in evolution. Trends Ecol. Evol. S0169534720301166.

Lenormand, T., L. M. Chevin, and T. Bataillon. 2016. Parallel evolution: what does it (not) tell us and why is it (still) interesting? P. *in* Chance in Evolution. Chicago: University Chicago Press.

Lenormand, T., D. Roze, and F. Rousset. 2009. Stochasticity in evolution. Trends Ecol. Evol. 24:157–165.

Li, H. 2018. Minimap2: pairwise alignment for nucleotide sequences. Bioinformatics 34:3094–3100.

Liu, H. 2014. Developing genomic resources for an emerging ecological model species *Senecio lautus*. PhD Thesis. The University of Queensland. doi: 10.14264/uql.2015.456.

Losos, J. B. 2011. Convergence, adaptation, and constraint. Evolution 65:1827–1840.

Lowry, D. B., S. Hoban, J. L. Kelley, K. E. Lotterhos, L. K. Reed, M. F. Antolin, and A. Storfer. 2017. Breaking RAD: an evaluation of the utility of restriction site-associated DNA sequencing for genome scans of adaptation. Mol. Ecol. Resour. 17:142–152.

Lowry, D. B., and J. H. Willis. 2010. A widespread chromosomal inversion polymorphism contributes to a major life-history transition, local adaptation, and reproductive isolation. PLoS Biol. 8:e1000500.

Mantel, N. 1967. The detection of disease clustering and a generalized regression approach. Cancer Res. 27:209–220.

Melo, M. C., A. Grealy, B. Brittain, G. M. Walter, and D. Ortiz-Barrientos. 2014. Strong extrinsic reproductive isolation between parapatric populations of an Australian groundsel. New Phytol. 203:323–334.

Melo, M. C., M. E. James, F. Roda, D. Bernal-Franco, M. J. Wilkinson, H. Liu, G. M. Walter, and D. Ortiz-Barrientos. 2019. Evidence for mutation-order speciation in an Australian wildflower. Preprint at https://doi.org/10.1101/692673.

Nelson, T. C., P. J. Monnahan, M. K. McIntosh, K. Anderson, E. MacArthur-Waltz, F. R. Finseth, J. K. Kelly, and L. Fishman. 2019. Extreme copy number variation at a TRNA ligase gene affecting phenology and fitness in yellow monkeyflowers. Mol. Ecol. 28:1460–1475.

Nosil, P., B. J. Crespi, and C. P. Sandoval. 2002. Host-plant adaptation drives the parallel evolution of reproductive isolation. Nature 417:440–443.

Ord, T. J., and T. C. Summers. 2015. Repeated evolution and the impact of evolutionary history on adaptation. BMC Evol. Biol. 15:137.

Perreault-Payette, A., A. M. Muir, F. Goetz, C. Perrier, E. Normandeau, P. Sirois, and L. Bernatchez. 2017. Investigating the extent of parallelism in morphological and genomic divergence among lake trout ecotypes in Lake Superior. Mol. Ecol. 26:1477–1497.

Pruitt, K. D. 2004. NCBI Reference Sequence (RefSeq): a curated non-redundant sequence database of genomes, transcripts and proteins. Nucleic Acids Res. 33:D501–D504.

Purcell, S., B. Neale, K. Todd-Brown, L. Thomas, M. A. R. Ferreira, D. Bender, J. Maller, P. Sklar, P. I. W. de Bakker, M. J. Daly, and P. C. Sham. 2007. PLINK: A tool set for whole-genome association and population-based linkage analyses. Am. J. Hum. Genet. 81:559–575.

R Core Team. 2017. R: A language and environment for statistical computing. R Foundation for Statistical Computing. Vienna, Austria. https://www.R-project.org/.

Richards, T. J., and D. Ortiz-Barrientos. 2016. Immigrant inviability produces a strong barrier to gene flow between parapatric ecotypes of *Senecio lautus*. Evolution 70:1239–1248.

Richards, T. J., G. M. Walter, K. McGuigan, and D. Ortiz-Barrientos. 2016. Divergent natural selection drives the evolution of reproductive isolation in an Australian wildflower. Evolution 70:1993–2003.

Rivas, M. J., M. Saura, A. Pérez-Figueroa, M. Panova, T. Johansson, C. André, A. Caballero, E. Rolán-Alvarez, K. Johannesson, and H. Quesada. 2018. Population genomics of parallel evolution in gene expression and gene sequence during ecological adaptation. Sci. Rep. 8:16147.

Roda, F., L. Ambrose, G. M. Walter, H. L. Liu, A. Schaul, A. Lowe, P. B. Pelser, P. Prentis, L. H. Rieseberg, and D. Ortiz-Barrientos. 2013a. Genomic evidence for the parallel evolution of coastal forms in the *Senecio lautus* complex. Mol Ecol 22:2941–2952.

Roda, F., H. Liu, M. J. Wilkinson, G. M. Walter, M. E. James, D. M. Bernal, M. C. Melo, A. Lowe, L. H. Rieseberg, P. Prentis, and D. Ortiz-Barrientos. 2013b. Convergence and divergence during the adaptation to similar environments by an Australian groundsel. Evolution 67:2515–2529.

Rosenblum, E. B., C. E. Parent, and E. E. Brandt. 2014. The molecular basis of phenotypic convergence. Annu. Rev. Ecol. Evol. Syst. 45:203–226.

Schluter, D. 2000. The ecology of adaptive radiation. Oxford: Oxford University Press.

Schluter, D., and L. M. Nagel. 1995. Parallel speciation by natural selection. Am. Nat. 146:292–301.

Schneider, C. A., W. S. Rasband, and K. W. Eliceiri. 2012. NIH Image to ImageJ: 25 years of image analysis. Nat. Methods 9:671–675.

Schrader, L., and J. Schmitz. 2019. The impact of transposable elements in adaptive evolution. Mol. Ecol. 28:1537–1549.

Schrider, D. R., M. W. Hahn, and D. J. Begun. 2016. Parallel evolution of copy-number variation across continents in *Drosophila melanogaster*. Mol. Biol. Evol. 33:1308–1316.

Smith, S. D., and M. D. Rausher. 2011. Gene loss and parallel evolution contribute to species difference in flower color. Mol. Biol. Evol. 28:2799–2810.

Soria-Carrasco, V., Z. Gompert, A. A. Comeault, T. E. Farkas, T. L. Parchman, J. S. Johnston, C. A. Buerkle, J. L. Feder, J. Bast, T. Schwander, S. P. Egan, B. J. Crespi, and P. Nosil. 2014. Stick insect genomes reveal natural selection’s role in parallel speciation. Science 344:738–742.

Staswick, P. E., B. Serban, M. Rowe, I. Tiryaki, M. T. Maldonado, M. C. Maldonado, and W. Suza. 2005. Characterization of an *Arabidopsis* enzyme family that conjugates amino acids to indole-3-acetic acid. Plant Cell 17:616–627.

Stern, D. L. 2013. The genetic causes of convergent evolution. Nat. Rev. Genet. 14:751–764.

Stern, D. L., and V. Orgogozo. 2009. Is genetic evolution predictable? Science 323:746–751.

Strohm, A. K., K. L. Baldwin, and P. H. Masson. 2012. Multiple roles for membrane-associated protein trafficking and signaling in gravitropism. Front. Plant Sci. 3.

Stuart, Y. E. 2019. Divergent uses of “parallel evolution” during the history of *The American Naturalist*. Am. Nat. 193:11–19.

Stuart, Y. E., T. Veen, J. N. Weber, D. Hanson, M. Ravinet, B. K. Lohman, C. J. Thompson, T. Tasneem, A. Doggett, R. Izen, N. Ahmed, R. D. H. Barrett, A. P. Hendry, C. L. Peichel, and D. I. Bolnick. 2017. Contrasting effects of environment and genetics generate a continuum of parallel evolution. Nat. Ecol. Evol. 1:0158.

Tenaillon, O., A. Rodriguez-Verdugo, R. L. Gaut, P. McDonald, A. F. Bennett, A. D. Long, and B. S. Gaut. 2012. The molecular diversity of adaptive convergence. Science 335:457–461.

Tiffin, P., and J. Ross-Ibarra. 2014. Advances and limits of using population genetics to understand local adaptation. Trends Ecol. Evol. 29:673–680.

Trucchi, E., B. Frajman, T. H. A. Haverkamp, P. Schönswetter, and O. Paun. 2017. Genomic analyses suggest parallel ecological divergence in *Heliosperma pusillum* (Caryophyllaceae). New Phytol. 216:267–278.

VanWallendael, A., A. Soltani, N. C. Emery, M. M. Peixoto, J. Olsen, and D. B. Lowry. 2019. A molecular view of plant local adaptation: Incorporating stress-response networks. Annu. Rev. Plant Biol. 70:559–583.

Verta, J. P., and F. C. Jones. 2019. Predominance of cis-regulatory changes in parallel expression divergence of sticklebacks. eLife 8:e43785.

Walter, G. M., J. D. Aguirre, M. W. Blows, and D. Ortiz-Barrientos. 2018a. Evolution of genetic variance during adaptive radiation. Am. Nat. 191:E108–E128.

Walter, G. M., M. J. Wilkinson, J. D. Aguirre, M. W. Blows, and D. Ortiz-Barrientos. 2018b. Environmentally induced development costs underlie fitness tradeoffs. Ecology 99:1391–1401.

Walter, G. M., M. J. Wilkinson, M. E. James, T. J. Richards, J. D. Aguirre, and D. Ortiz-Barrientos. 2016. Diversification across a heterogeneous landscape. Evolution 70:1979–1992.

Wilkinson, M. J., F. Roda, G. M. Walter, M. E. James, R. Nipper, J. Walsh, H. L. North, C. A. Beveridge, and D. Ortiz-Barrientos. 2021. Adaptive divergence in shoot gravitropism creates hybrid sterility in an Australian wildflower. Preprint at https://doi.org/10.1101/845354.

Wood, T. E., J. M. Burke, and L. H. Rieseberg. 2005. Parallel genotypic adaptation: when evolution repeats itself. Genetica 123:157–170.

Yeaman, S. 2015. Local adaptation by alleles of small effect. Am. Nat. 186:S74–S89.

