## Supplementary Figures and Tables for "Phenotypic and genotypic parallel evolution in parapatric ecotypes of *Senecio*"

### Supporting information

#### Table S1. *Senecio lautus* sampling locations in this study

Sampling locations of the 22 *Senecio lautus* Dune and Headland populations. Coordinates represent the mid-point of each population. N genetics and N phenotype represent the sample sizes used for the genotypic and phenotypic analysis respectively. Note: N genetics corresponds to the final number of individuals after removing those with low coverage. Pairs in bold are sister-taxa within the phylogeny.

| Clade | Population code | Location | Ecotype | Pair | Coordinates | N genetics | N phenotype |
|---|---|---|---|---|---|---|---|
| Eastern | D00 | QLD: Stradbroke Island | Dune | D00-H00 | S27° 31.153' E153° 30.189' | 62 | 30 |
| Eastern | H00 | QLD: Stradbroke Island | Headland | D00-H00 | S27° 26.140' E153° 32.749' | 63 | 30 |
| Eastern | D02 | QLD: Southport | Dune | D02-H04 | S27° 56.846' E153° 25.736' | 62 | 30 |
| Eastern | H04 | NSW: Byron Bay | Headland | D02-H04 | S28° 38.060' E153° 38.268' | 62 | 30 |
| Eastern | D03 | NSW: Cabarita | Dune | **D03-H02** | S28° 19.794' E153° 34.264' | 61 | 30 |
| Eastern | H02 | NSW: Cabarita | Headland | **D03-H02** | S28° 21.013' E153° 34.676' | 61 | 30 |
| Eastern | D01 | NSW: Lennox Head | Dune | D01-H01 | S28° 46.858' E153° 35.655' | 60 | 30 |
| Eastern | H01 | NSW: Lennox Head | Headland | D01-H01 | S28° 48.813' E153° 36.313' | 58 | 30 |
| Eastern | D04 | NSW: Coffs Harbour | Dune | **D04-H05** | S30° 18.946' E153° 08.142' | 62 | 30 |
| Eastern | H05 | NSW: Coffs Harbour | Headland | **D04-H05** | S30° 18.741' E153° 08.676' | 62 | 30 |
| Eastern | D05 | NSW: South West Rocks | Dune | **D05-H06** | S30° 53.027' E153° 04.037' | 62 | 30 |
| Eastern | H06 | NSW: South West Rocks | Headland | **D05-H06** | S30° 52.710' E153° 04.549' | 62 | 30 |
| Southern | H07 | NSW: Port Macquarie | Headland | - | S31° 28.526' E152° 56.219 | - | 30 |
| Southern | H03 | NSW: Kiama | Headland | - | S34° 40.301' E150° 51.704' | - | 29 |
| Southern | D12 | NSW: Bermagui | Dune | D12-H14 | S36° 28.346' E150° 03.581' | 62 | 31 |
| Southern | H14 | NSW: Green Cape | Headland | D12-H14 | S37° 15.748' E150° 02.991' | 62 | 30 |
| Southern | D32 | VIC: Cape Bridgewater | Dune | **D32-H12** | S38° 19.631' E141° 23.772' | 62 | 30 |
| Southern | H12 | VIC: Cape Bridgewater | Headland | **D32-H12** | S38° 22.728' E141° 22.018' | 63 | 30 |
| Southern | D14 | TAS: Port Arthur | Dune | **D14-H15** | S43° 10.550' E147° 51.267' | 12 | - |
| Southern | H15 | TAS: Port Arthur | Headland | **D14-H15** | S43° 11.240' E147° 50.672' | 11 | - |
| Western | D35 | WA: Isthmus Hill | Dune | - | S35° 05.885' E117° 59.182' | - | 34 |
| Western | D09 | WA: Leeuwin-Naturaliste National Park | Dune | - | S33° 46.239' E114° 59.541' | - | 31 |

**Table S2.** ***Senecio lautus*** **phenotypic traits**

Traits with asterisks were removed from the analysis due to high correlations (> 0.8).

| Trait category | Trait | Description |
|---|---|---|
| Plant architecture | Vegetative height | Base of plant to highest vegetative leaf (not including flowers) |
| | Widest width | Widest with of plant between vegetative leaves |
| | Narrowest width* | Narrowest width of plant between vegetative leaves |
| | Main stem angle | Angle of main stem measured from base of soil |
| | Main stem diameter | Diameter of main stem measured 1cm above soil |
| | Primary branch angle† | Average angle of primary branches |
| Leaf | Leaf | Leaf area |
| | Perimeter* | Leaf perimeter |
| | Width* | Leaf width |
| | Height* | Leaf height |
| | Elongation | Length to width ratio |
| | Compactness* | Squared perimeter to area ratio |
| | Dissection | Perimeter to length ratio |
| | Circularity | $4\pi$ (area/perimeter$^2$) |

† We note that in previous *Senecio lautus* papers before 2021, this was referred to as 'secondary branch angle'

**Table S3. Pairwise change in lengths (ΔL) and angles (θ) for the removal of single traits in coastal ecotypes of *S. lautus***

For each matrix, values below the diagonal represent the either the change in lengths or the angles (in degrees), above the diagonal are the P-values. Shaded cells donate parallel pairs ($P > 0.01$, i.e. $\theta \approx 0°$). See Supplementary excel document for Table S3.

### Table S4. Demographic and environmental data for *S. lautus* population pairs

Rates of gene flow (*m*, backward in time) and divergence times were calculated using the same dataset in *fastsimcoal2* (see James et al. 2021 for details). *m* (D -> H): gene flow from the Dune to the Headland. 2N*m* (H -> D): gene flow Headland to the Dune. *m* (average): average gene flow. See (Roda et al. 2013) for details of the environmental distances.

| **Pair** | ***m*** **(D -> H)** | ***m*** **(H -> D)** | ***m*** **(average)** | **Divergence time (years)** | **Environmental distance** |
|---|---|---|---|---|---|
| D00-H00 | 1.18E-05 | 3.24E-06 | 7.53E-06 | 71,945 | 3.23 |
| D02-H04 | NA | NA | NA | NA | NA |
| D03-H02 | 2.73E-05 | 2.08E-06 | 1.47E-05 | 44,190 | 10.26 |
| D01-H01 | 8.13E-06 | 7.85E-06 | 7.99E-06 | 71,918 | 13.68 |
| D04-H05 | 2.96E-05 | 3.69E-05 | 3.33E-05 | 37,211 | 6.31 |
| D05-H06 | 6.56E-06 | 1.14E-05 | 8.99E-06 | 67,927 | NA |
| D12-H14 | 1.69E-06 | 4.24E-06 | 2.97E-06 | 110,018 | 5.14 |
| D32-H12 | 1.59E-05 | 5.25E-05 | 3.42E-05 | 38,706 | NA |

**Table S5. Linear discriminant analysis of all traits between Dune and Headland ecotypes**

| Trait Category | Trait | LDA |
|---|---|---|
| Plant architecture | Vegetative height | -0.26 |
| | Widest width | 0.41 |
| | Main stem angle | -0.15 |
| | Main stem diameter | -0.36 |
| | Primary branch angle | -0.61 |
| Leaf | Area | -0.78 |
| | Elongation | -0.01 |
| | Dissection | 0.51 |
| | Circularity | 0.45 |

**Table S6. Trait-by-trait linear models and partial effect sizes**

F-values and P-values for the ecotype (Ecotype $F_{1,465}$; Ecotype P-value) and interaction (Ecotype x Pair $F_{7,465}$; Ecotype × Pair P-value) for trait-by-trait linear models (*trait* = *ecotype* + *pair* + *ecotype* × *pair*). Partial effect sizes (partial $\eta^2$) for the Ecotype, Pair and Interaction (Ecotype × Pair) for each trait.

| Trait Category | Trait | Ecotype $F_{1,465}$ | Ecotype P-value | Ecotype × Pair $F_{7,465}$ | Ecotype × Pair P-value | Ecotype partial $\eta^2$ | Pair partial $\eta^2$ | Ecotype × Pair partial $\eta^2$ |
|---|---|---|---|---|---|---|---|---|
| Plant architecture | Vegetative height | 962.65 | $< 2.2 \times 10^{-16}$ *** | 17.63 | $< 2.2 \times 10^{-16}$ *** | 0.67 | 0.26 | 0.21 |
| | Widest width | 102.62 | $< 2.2 \times 10^{-16}$ *** | 8.14 | $2.3 \times 10^{-9}$ *** | 0.18 | 0.30 | 0.11 |
| | Main stem angle | 111.35 | $< 2.2 \times 10^{-16}$ *** | 7.20 | $3.4 \times 10^{-8}$ *** | 0.19 | 0.06 | 0.10 |
| | Main stem diameter | 133.75 | $< 2.2 \times 10^{-16}$ *** | 11.26 | $3.3 \times 10^{-13}$ *** | 0.22 | 0.07 | 0.14 |
| | Primary branch angle | 566.05 | $< 2.2 \times 10^{-16}$ *** | 2.39 | 0.02092 * | 0.55 | 0.08 | 0.03 |
| Leaf | Area | 889.52 | $< 2.2 \times 10^{-16}$ *** | 16.30 | $< 2.2 \times 10^{-16}$ *** | 0.66 | 0.26 | 0.20 |
| | Elongation | 398.11 | $< 2.2 \times 10^{-16}$ *** | 19.79 | $< 2.2 \times 10^{-16}$ *** | 0.46 | 0.26 | 0.23 |
| | Dissection | 84.64 | $< 2.2 \times 10^{-16}$ *** | 20.92 | $< 2.2 \times 10^{-16}$ *** | 0.15 | 0.25 | 0.24 |
| | Circularity | 356.68 | $< 2.2 \times 10^{-16}$ *** | 45.92 | $< 2.2 \times 10^{-16}$ *** | 0.43 | 0.46 | 0.41 |

**Table S7. Pairwise change in lengths (ΔL) for all traits in coastal ecotypes of *S. lautus***

Values below the diagonal represent the change in lengths, above the diagonal are the P-values. Shaded cells donate parallel pairs (P > 0.01, i.e. ΔL ≈ 0°).

| | D00-H00 | D01-H01 | D02-H04 | D03-H02 | D04-H05 | D05-H06 | D12-H14 | D32-H12 |
|---|---|---|---|---|---|---|---|---|
| **D00-H00** | - | 0.3239 | 0.0001 | 0.0001 | 0.0001 | 0.4949 | 0.0001 | 0.0001 |
| **D01-H01** | 0.27 | - | 0.0001 | 0.0001 | 0.0001 | 0.0763 | 0.0001 | 0.0001 |
| **D02-H04** | 2.45 | 2.72 | - | 0.0548 | 0.0002 | 0.0001 | 0.0001 | 0.5683 |
| **D03-H02** | 1.86 | 2.13 | 0.59 | - | 0.0435 | 0.0001 | 0.0367 | 0.1627 |
| **D04-H05** | 1.27 | 1.54 | 1.18 | 0.59 | - | 0.0003 | 0.8839 | 0.0003 |
| **D05-H06** | 0.20 | 0.47 | 2.26 | 1.66 | 1.08 | - | 0.0007 | 0.0001 |
| **D12-H14** | 1.24 | 1.51 | 1.22 | 0.62 | 0.04 | 1.04 | - | 0.0005 |
| **D32-H12** | 2.29 | 2.56 | 0.16 | 0.43 | 1.02 | 2.10 | 1.05 | - |

**Table S8. Pairwise change in lengths (ΔL) for plant architecture traits in coastal ecotypes of *S. lautus***

Values below the diagonal represent the change in lengths, above the diagonal are the P-values. Shaded cells donate parallel pairs (P > 0.01, i.e. ΔL ≈ 0°).

| | D00-H00 | D01-H01 | D02-H04 | D03-H02 | D04-H05 | D05-H06 | D12-H14 | D32-H12 |
|---|---|---|---|---|---|---|---|---|
| **D00-H00** | - | 0.1366 | 0.0001 | 0.0001 | 0.0001 | 0.6428 | 0.0016 | 0.0008 |
| **D01-H01** | 0.44 | - | 0.0001 | 0.0001 | 0.0001 | 0.0394 | 0.0002 | 0.0001 |
| **D02-H04** | 1.55 | 1.99 | - | 0.0926 | 0.4009 | 0.0001 | 0.0477 | 0.1153 |
| **D03-H02** | 2.08 | 2.52 | 0.53 | - | 0.0056 | 0.0001 | 0.0001 | 0.0017 |
| **D04-H05** | 1.32 | 1.76 | 0.23 | 0.77 | - | 0.0001 | 0.1633 | 0.3648 |
| **D05-H06** | 0.14 | 0.58 | 1.41 | 1.94 | 1.18 | - | 0.0070 | 0.0032 |
| **D12-H14** | 0.94 | 1.38 | 0.61 | 1.14 | 0.37 | 0.80 | - | 0.6626 |
| **D32-H12** | 1.07 | 1.52 | 0.48 | 1.01 | 0.24 | 0.94 | 0.13 | - |

**Table S9. Pairwise change in lengths (ΔL) for leaf traits in coastal ecotypes of *S. lautus***

Values below the diagonal represent the change in lengths, above the diagonal are the P-values. Shaded cells donate parallel pairs (P > 0.01, i.e. ΔL ≈ 0°).

| | D00-H00 | D01-H01 | D02-H04 | D03-H02 | D04-H05 | D05-H06 | D12-H14 | D32-H12 |
|---|---|---|---|---|---|---|---|---|
| **D00-H00** | - | 0.9650 | 0.0001 | 0.0740 | 0.0239 | 0.4917 | 0.0006 | 0.0001 |
| **D01-H01** | 0.01 | - | 0.0001 | 0.1268 | 0.0509 | 0.5771 | 0.0046 | 0.0001 |
| **D02-H04** | 1.90 | 1.89 | - | 0.0001 | 0.0001 | 0.0001 | 0.0015 | 0.4492 |
| **D03-H02** | 0.34 | 0.34 | 1.56 | - | 0.6506 | 0.3263 | 0.0352 | 0.0001 |
| **D04-H05** | 0.43 | 0.42 | 1.47 | 0.09 | - | 0.1646 | 0.0674 | 0.0001 |
| **D05-H06** | 0.14 | 0.13 | 1.77 | 0.21 | 0.30 | - | 0.0052 | 0.0001 |
| **D12-H14** | 0.81 | 0.80 | 1.10 | 0.46 | 0.37 | 0.67 | - | 0.0001 |
| **D32-H12** | 2.08 | 2.07 | 0.17 | 1.73 | 1.64 | 1.94 | 1.27 | - |

**Table S10. Pairwise angles (θ) for all traits in coastal ecotypes of *S. lautus***

Values below the diagonal represent the angles (in degrees), above the diagonal are the P-values. Shaded cells donate parallel pairs (P > 0.01, i.e. θ ≈ 0°).

| | D00-H00 | D01-H01 | D02-H04 | D03-H02 | D04-H05 | D05-H06 | D12-H14 | D32-H12 |
|---|---|---|---|---|---|---|---|---|
| **D00-H00** | - | 0.0001 | 0.0001 | 0.0001 | 0.0001 | 0.0001 | 0.0001 | 0.0001 |
| **D01-H01** | 43.38 | - | 0.0001 | 0.0001 | 0.0001 | 0.0001 | 0.0018 | 0.0001 |
| **D02-H04** | 43.73 | 49.83 | - | 0.0001 | 0.0001 | 0.0001 | 0.0001 | 0.0001 |
| **D03-H02** | 31.00 | 50.52 | 40.33 | - | 0.0105 | 0.0001 | 0.0001 | 0.0001 |
| **D04-H05** | 26.57 | 42.70 | 42.93 | 14.51 | - | 0.0001 | 0.0001 | 0.0001 |
| **D05-H06** | 61.69 | 49.81 | 62.81 | 39.29 | 39.30 | - | 0.0001 | 0.0001 |
| **D12-H14** | 50.10 | 35.99 | 34.81 | 34.08 | 35.69 | 35.34 | - | 0.0010 |
| **D32-H12** | 50.27 | 34.82 | 45.36 | 33.59 | 33.74 | 24.43 | 20.14 | - |

**Table S11. Pairwise angles (θ) for plant architecture traits in coastal ecotypes of *S. lautus***

Values below the diagonal represent the angles (in degrees), above the diagonal are the P-values. Shaded cells donate parallel pairs ($P > 0.01$, i.e. $\theta \approx 0°$).

| | D00-H00 | D01-H01 | D02-H04 | D03-H02 | D04-H05 | D05-H06 | D12-H14 | D32-H12 |
|---|---|---|---|---|---|---|---|---|
| **D00-H00** | - | 0.0021 | 0.0025 | 0.0049 | 0.0022 | 0.0003 | 0.0013 | 0.0005 |
| **D01-H01** | 49.70 | - | 0.0001 | 0.0001 | 0.0001 | 0.0005 | 0.0002 | 0.0001 |
| **D02-H04** | 26.78 | 62.07 | - | 0.2492 | 0.0270 | 0.0002 | 0.0002 | 0.0161 |
| **D03-H02** | 23.10 | 60.56 | 9.27 | - | 0.0395 | 0.0001 | 0.0009 | 0.0043 |
| **D04-H05** | 27.06 | 50.46 | 15.63 | 13.48 | - | 0.0198 | 0.1135 | 0.5276 |
| **D05-H06** | 43.11 | 50.21 | 30.17 | 30.37 | 18.95 | - | 0.0051 | 0.0835 |
| **D12-H14** | 31.30 | 44.17 | 26.61 | 22.66 | 13.08 | 23.40 | - | 0.1199 |
| **D32-H12** | 33.60 | 50.15 | 18.37 | 19.47 | 8.49 | 17.08 | 14.04 | - |

**Table S12. Pairwise angles (θ) for leaf traits in coastal ecotypes of *S. lautus***

Values below the diagonal represent the angles (in degrees), above the diagonal are the P-values. Shaded cells donate parallel pairs ($P > 0.01$, i.e. $\theta \approx 0°$).

| | D00-H00 | D01-H01 | D02-H04 | D03-H02 | D04-H05 | D05-H06 | D12-H14 | D32-H12 |
|---|---|---|---|---|---|---|---|---|
| **D00-H00** | - | 0.0014 | 0.0001 | 0.0001 | 0.0012 | 0.0001 | 0.0001 | 0.0001 |
| **D01-H01** | 37.73 | - | 0.0001 | 0.1043 | 0.0172 | 0.0001 | 0.0531 | 0.0009 |
| **D02-H04** | 54.25 | 40.54 | - | 0.0001 | 0.0001 | 0.0001 | 0.0001 | 0.0001 |
| **D03-H02** | 31.56 | 19.33 | 58.04 | - | 0.0907 | 0.0001 | 0.0002 | 0.0001 |
| **D04-H05** | 21.88 | 26.54 | 60.06 | 11.42 | - | 0.0001 | 0.0001 | 0.0001 |
| **D05-H06** | 75.54 | 48.76 | 82.96 | 44.26 | 54.12 | - | 0.0001 | 0.0001 |
| **D12-H14** | 63.73 | 26.53 | 40.49 | 42.27 | 51.98 | 43.90 | - | 0.0060 |
| **D32-H12** | 58.83 | 24.12 | 56.47 | 29.60 | 40.89 | 26.79 | 20.31 | - |

**Table S13. Shared outlier nucleotide sites between replicate population pairs from coastal ecotypes of *S. lautus***

Values below the diagonal represent the number of shared outlier nucleotide sites. Values above the diagonal are the P-values of the probability that the common nucleotide sites are shared by chance, calculated from the hypergeometric distribution.

| | D00-H00 | D01-H01 | D02-H04 | D03-H02 | D04-H05 | D05-H06 | D12-H14 | D14-H15 | D32-H12 |
|---|---|---|---|---|---|---|---|---|---|
| **D00-H00** | - | 3.75E-58 | 7.72E-54 | 3.98E-37 | 5.31E-51 | 8.48E-42 | 8.13E-50 | 2.74E-07 | 1.68E-32 |
| **D01-H01** | 169 | - | 3.59E-58 | 9.08E-52 | 1.14E-62 | 1.52E-48 | 1.39E-51 | 4.07E-05 | 6.49E-33 |
| **D02-H04** | 189 | 161 | - | 1.58E-49 | 3.52E-32 | 1.10E-53 | 1.20E-37 | 1.80E-03 | 3.40E-44 |
| **D03-H02** | 178 | 164 | 187 | - | 8.27E-46 | 1.36E-46 | 3.76E-44 | 2.63E-05 | 6.47E-36 |
| **D04-H05** | 195 | 174 | 155 | 192 | - | 5.31E-51 | 3.60E-44 | 1.07E-05 | 4.69E-34 |
| **D05-H06** | 165 | 144 | 173 | 176 | 178 | - | 2.27E-58 | 1.15E-11 | 7.98E-49 |
| **D12-H14** | 226 | 185 | 193 | 222 | 216 | 218 | - | 6.91E-10 | 7.40E-68 |
| **D14-H15** | 31 | 21 | 21 | 28 | 28 | 35 | 41 | - | 1.13E-03 |
| **D32-H12** | 207 | 164 | 215 | 220 | 210 | 214 | 311 | 30 | - |

**Table S14.** ***S. lautus*** **parallel gene functions**

Summary of the parallel gene functions detected in either *Approach 1*, *2* or *3*. Gene is the gene name. Gene ontology is the *DAVID GOTERM_BP_DIRECT* (denoted below with *BP*), where absent, the *GOTERM_CC_DIRECT* was used (*CC*), and where absent, the *GOTERM_MF_DIRECT* (*MF*) was used. *Approach 1* denotes if the outlier was detected due to high differentiation between ecotypes. *Approach 2* denotes if the outlier was detected due to high differentiation at each locality (note: genes only shown if they were highly differentiated in all nine, or eight replicate pairs). *Approach 3* denotes outliers that contain concordant allele frequency changes across replicate pairs. See Supplementary Methods S1 for extra details. See Supplementary excel document for Table S14.

**Table S15. Shared outlier genes between replicate population pairs from coastal ecotypes of *S. lautus***

Values below the diagonal represent the number of shared outlier genes. Values above the diagonal are the P-values of the probability that the common genes are shared by chance, calculated from the hypergeometric distribution.

| | D00-H00 | D01-H01 | D02-H04 | D03-H02 | D04-H05 | D05-H06 | D12-H14 | D14-H15 | D32-H12 |
|---|---|---|---|---|---|---|---|---|---|
| **D00-H00** | - | 2.10E-40 | 2.76E-53 | 1.54E-34 | 6.83E-33 | 2.47E-39 | 9.75E-24 | 2.05E-05 | 1.01E-19 |
| **D01-H01** | 123 | - | 1.13E-52 | 1.18E-44 | 5.03E-45 | 2.07E-35 | 2.88E-29 | 7.56E-04 | 1.79E-22 |
| **D02-H04** | 156 | 128 | - | 8.25E-47 | 2.06E-34 | 1.26E-42 | 5.36E-31 | 3.48E-05 | 1.15E-34 |
| **D03-H02** | 137 | 122 | 142 | - | 1.11E-34 | 1.78E-35 | 2.78E-22 | 7.95E-05 | 1.16E-24 |
| **D04-H05** | 141 | 127 | 133 | 136 | - | 6.85E-34 | 1.52E-20 | 2.30E-03 | 7.76E-20 |
| **D05-H06** | 146 | 114 | 140 | 134 | 138 | - | 1.32E-32 | 1.25E-08 | 5.61E-24 |
| **D12-H14** | 171 | 140 | 168 | 158 | 164 | 179 | - | 1.50E-03 | 2.34E-17 |
| **D14-H15** | 32 | 22 | 29 | 29 | 27 | 37 | 40 | - | 0.10091 |
| **D32-H12** | 168 | 133 | 177 | 166 | 167 | 170 | 231 | 35 | - |

**Table S16. Shared biological functions between replicate population pairs from coastal ecotypes of *S. lautus***

Summary of the 17 enriched biological functions across the 9 replicate pairs inferred in *DAVID*. P-values represent the EASE score, a modified Fisher Exact P-value. 'NA' denotes the function was not enriched within the pair.

| Biological function | D00-H00 | D02-H04 | D03-H02 | D01-H01 | D04-H05 | D05-H06 | D12-H14 | D14-H15 | D32-H12 |
|---|---|---|---|---|---|---|---|---|---|
| Chloroplast | 7.25E-04 | 4.91E-05 | 1.08E-04 | 6.63E-03 | 8.56E-03 | 3.02E-02 | 3.57E-10 | NA | 4.79E-05 |
| Nucleotide/ATP-binding | 1.12E-05 | 3.05E-03 | 7.00E-07 | 5.32E-04 | 7.74E-04 | 2.33E-02 | 7.87E-03 | NA | 3.77E-07 |
| Membrane | 8.50E-04 | NA | NA | 3.55E-05 | 1.22E-02 | 2.25E-02 | 1.67E-03 | NA | 9.48E-03 |
| Helicase | 2.38E-03 | NA | 7.20E-03 | NA | 2.69E-02 | 1.62E-02 | 8.08E-04 | NA | 3.90E-02 |
| DNA repair/DNA damage | NA | 3.81E-02 | 4.32E-02 | 7.01E-04 | 4.65E-02 | NA | 1.42E-03 | NA | 3.31E-02 |
| Flowering | NA | NA | 3.36E-04 | 5.95E-03 | NA | 3.60E-02 | NA | 2.79E-02 | 1.24E-02 |
| Protein-protein interactions | NA | NA | NA | 8.00E-03 | 1.90E-02 | 2.96E-02 | NA | 3.38E-02 | 1.32E-03 |
| Carbon metabolism | 2.37E-02 | NA | NA | 2.35E-02 | 4.40E-02 | NA | NA | 5.93E-03 | 2.14E-02 |
| Zinc finger | 2.47E-02 | 2.25E-02 | NA | NA | NA | 1.73E-02 | 2.28E-03 | NA | NA |
| Nucleus | NA | 2.59E-02 | 1.34E-02 | NA | NA | NA | 4.36E-03 | NA | 4.44E-03 |
| ATPase dependent activity | NA | NA | 4.08E-03 | NA | NA | 4.02E-03 | 4.20E-04 | 3.55E-02 | NA |
| Catalytic activity | NA | NA | 1.34E-02 | NA | NA | 6.62E-03 | 5.54E-04 | NA | NA |
| Stress response | NA | NA | NA | NA | NA | 3.18E-02 | 9.14E-03 | NA | NA |
| Auxin pathway | NA | NA | NA | NA | NA | 1.16E-02 | NA | NA | NA |
| Amino acid transport | NA | NA | NA | NA | NA | NA | NA | NA | 4.29E-03 |
| Glycoprotein | 1.63E-02 | NA | NA | NA | NA | NA | NA | NA | NA |
| Endoplasmic reticulum | NA | NA | NA | NA | NA | 3.80E-02 | NA | NA | NA |

**Table S17. Outlier genes within the *chloroplast* category**

Gene is the gene name. Gene ontology is the *DAVID GOTERM_BP_DIRECT* (denoted below with *BP*), and where absent, the *GOTERM_CC_DIRECT* was used (*CC*). Number of replicate pairs denotes how many localities the gene was detected as an outlier. See Supplementary excel document for Table S17.

**Table S18. Outlier genes within the *nucleotide/ATP-binding* category**

Gene is the gene name. Gene ontology is the *DAVID GOTERM_BP_DIRECT* (denoted below with *BP*), and where absent, the *GOTERM_CC_DIRECT* was used (*CC*). Number of replicate pairs denotes how many localities the gene was detected as an outlier. See Supplementary excel document for Table S18.

**Table S19. Auxin genes divergent between Dune and Headland *S. lautus* ecotypes in at least one locality**

Gene is the gene name. Gene ontology contains all ontologies from the *DAVID* analysis. Number of replicate pairs denotes how many localities the gene was detected as an outlier. See Supplementary excel document for Table S19.

**Figure S1. Frequency distribution of haploblocks less than 1000bp**

Haploblocks were calculated per population in *PLINK*. 95% of all haploblocks fall below 1000bp (those plotted in the figure), 91% fall below 300bp. Mean of all haploblocks is 359bp, median is 42bp. This reveals that linkage within each population decays quickly, and SNPs sequenced in different RAD-tags are likely unlinked. Therefore, within our work we can treat most SNPs as independent, unless the SNPs are in very close physical proximity. Although the presence of linked SNPs might slightly overestimate the extent of parallel evolution at the level of the nucleotide site, linkage between genes would generally be low, and we can treat separate genes as largely independent.

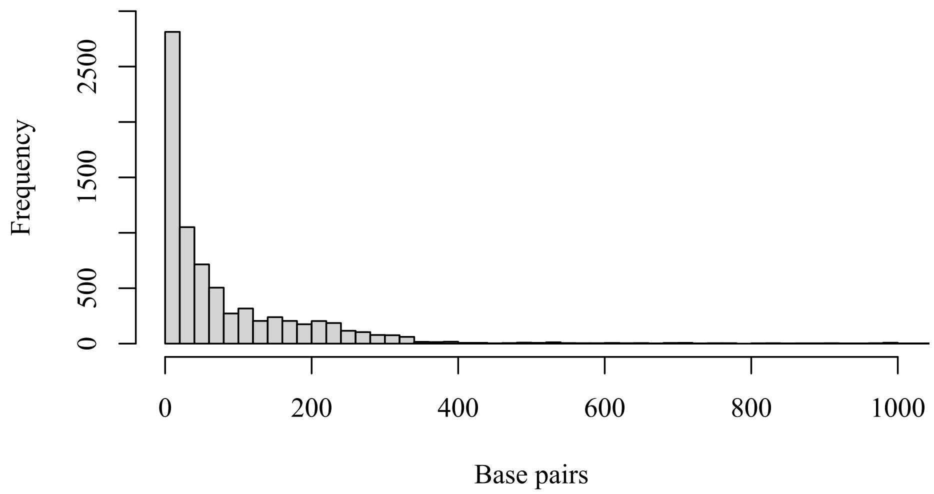

**Figure S2. Principal components analysis of plant architecture and leaf traits in coastal ecotypes of *S. lautus***

Principal component analysis of Dune (orange) and Headland (green) phenotypes for **(A)** five plant architecture, and **(B)** four leaf traits across 20 populations. Ecotypes are delimited by 70% probability ellipses.

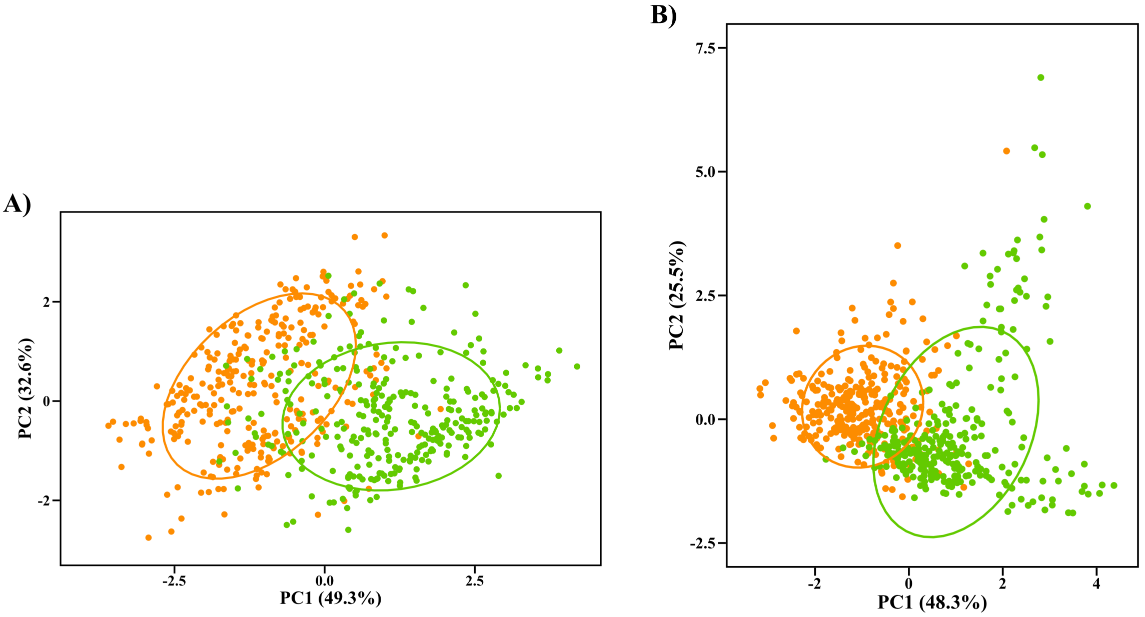

**Figure S3. Trait-by-trait effect sizes in coastal ecotypes of *S. lautus***

Partial effect sizes (partial $\eta^2$) for the ecotype and pair for the trait-by-trait linear models, each dot representing a single trait. The blue dot represents the partial effect size for all traits combined within the MANOVA. Dashed line is a 1:1 ratio, where points above the line represent a larger contribution of broad- to narrow-sense divergence. See Table S4 for details.

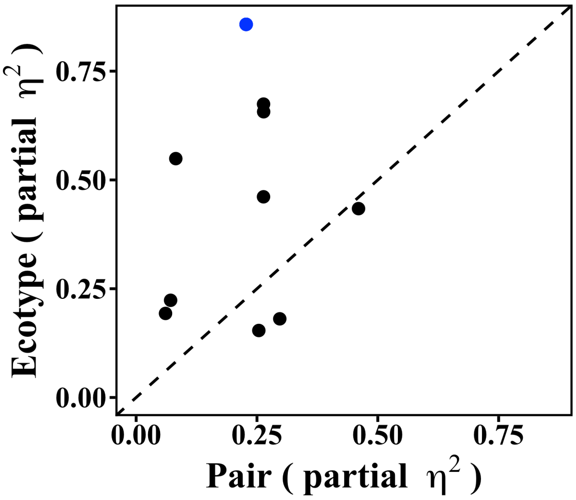

**Figure S4. Phenotypic Change Vector Analysis for plant architecture traits in coastal ecotypes from *S. lautus***

**(A)** PC1 and PC2 for five plant architecture traits across eight replicate Dune-Headland pairs. Each dot represents the population centroid (multivariate phenotypic mean), ± SE. The Dune (orange) and Headland (green) populations of a replicate pair are connected with a line. **(B)** Frequency distribution of the 28 pairwise phenotypic divergences (ΔL) between Dune-Headland replicate pairs (Supplementary Table S7). **(C)** Frequency distribution of the 28 pairwise contribution of traits (θ) between Dune-Headland replicate pairs (Supplementary Table S10).

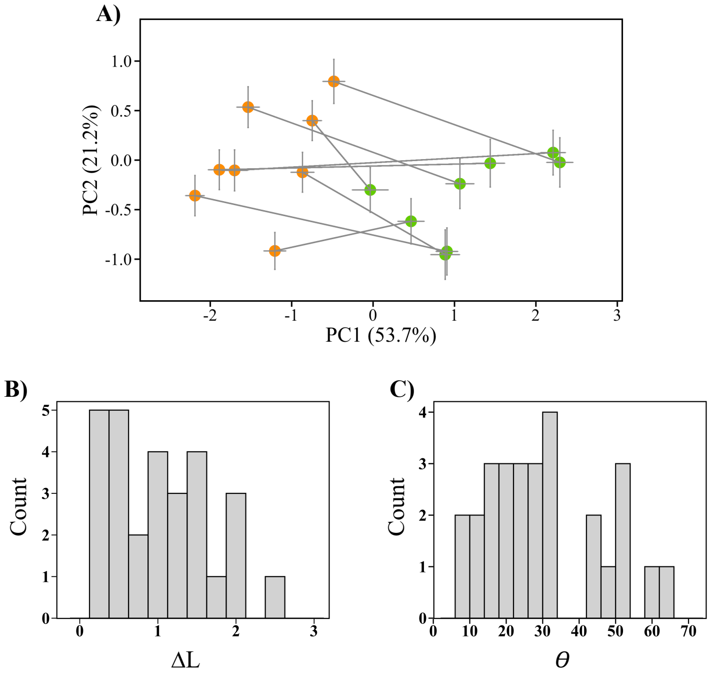

**Figure S5. Phenotypic Change Vector Analysis for leaf traits in costal ecotypes from *S. lautus***

**(A)** PC1 and PC2 for four leaf traits across eight replicate Dune-Headland pairs. Each dot represents the population centroid (multivariate phenotypic mean), ± SE. The Dune (orange) and Headland (green) populations of a replicate pair are connected with a line. **(B)** Frequency distribution of the 28 pairwise phenotypic divergences (ΔL) between Dune-Headland replicate pairs (Supplementary Table S8). **(C)** Frequency distribution of the 28 pairwise contribution of traits (θ) between Dune-Headland replicate pairs (Supplementary Table S11).

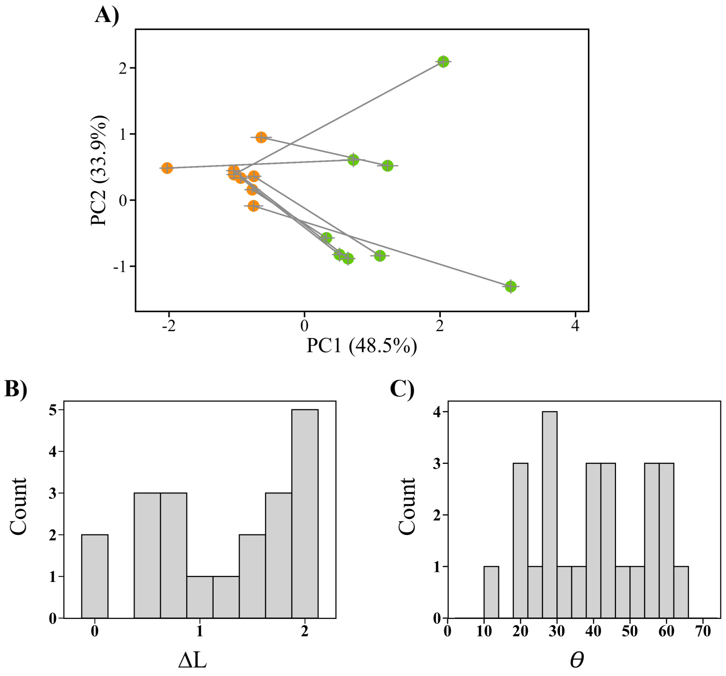

**Figure S6. Relative contributions of genotypic parallel and non-parallel evolution**

Partial effect sizes (partial $\eta^2$) for the ecotype and the pair from linear models for all sequenced nucleotide sites **(A)** and genes **(B)**. Each dot represents either a single SNP **(A)** or gene **(B)**. Most points fall below the dashed 1:1 ratio line, indicating that the variation in Dune-Headland divergence is largely unique to replicate pairs (non-parallel), rather than shared across localities (parallel). The blue dots denote the best candidates for parallel evolution (those in Figure 5A) at the level of the SNP **(A)** and gene **(B)**. The data in **(A)** and **(B)** is plotted as frequency distributions for the nucleotide sites **(C)** and genes **(D)**. Values represent the distance of the SNP or gene from the 1:1 dashed line of equal effect. Positive values indicate more parallel evolution, whereas negative values indicate more non-parallel evolution. As most values fall below zero, between-ecotype variation at the level of the nucleotide site and gene is mainly unique to replicate pairs.

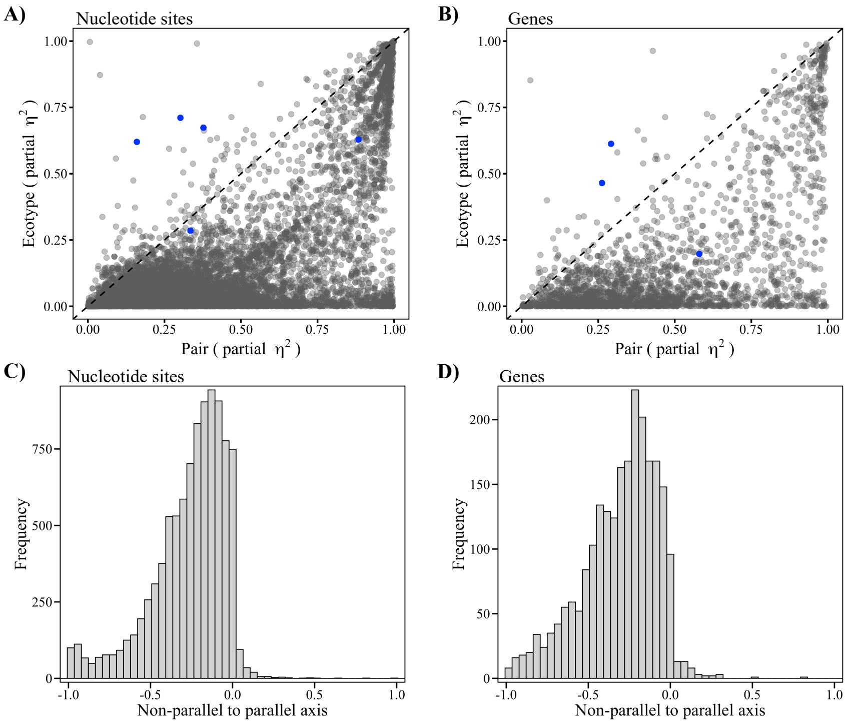

**Figure S7. Parallel nucleotide sites with concordant allele frequency differences**

Parallel nucleotide sites with concordant allele frequency differences (as detected in *Approach 3*) in either all nine (S-statistic = 9, P = 0.004), or eight (S-statistic = 8, P = 0.04) replicate pairs. Dots represent the allele frequency value (of the reference allele) for each population. Lines connect the Dune (orange) populations to their Headland (green) pair at each locality. Dashed lines represent pairs whose Dune-Headland change in allele frequency is in the opposite direction from the majority of pairs. *G* denotes SNPs that occur within genic regions. These genes are as follows (left to right, top to bottom): *AT5G14540*, NA, *AT5G04420*, *CNGC1*, *OVA9*, *AT5G65740*, *EMB3144*, *HCT*. Asterisks denote SNPs that were also detected as outliers within *Approach 1* (see *Methods* for details).

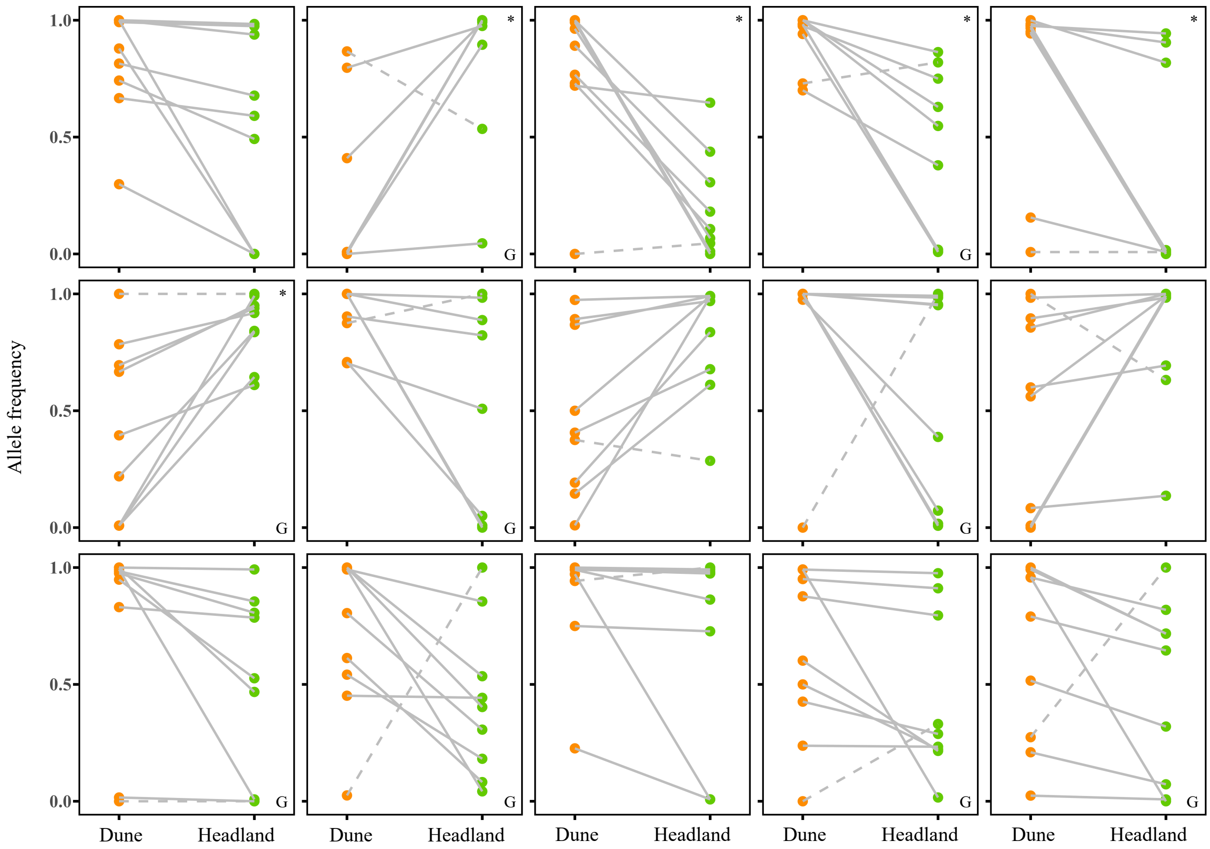

**Figure S8. Patterns of Dune-Headland FST, CSS and Δp in *S. lautus***

**(A)** Relationship between the change in allele frequency ($\Delta p$) and $F_{ST}$ comparing all Dune vs all Headland individuals, each datapoint representing a SNP. Blue denotes the top 1% $F_{ST}$ values. **(B)** Relationship of $\Delta p$ to the cluster separation score (CSS). Blue denotes the top 1% CSS values. **(C)** Relationship of $F_{ST}$ and CSS. Blue denotes the SNPs considered as outliers, being detected as outliers in at least two of the following approaches: top 1% $F_{ST}$, top 1% CSS, and *BayeScan* posterior probability > 0.91.

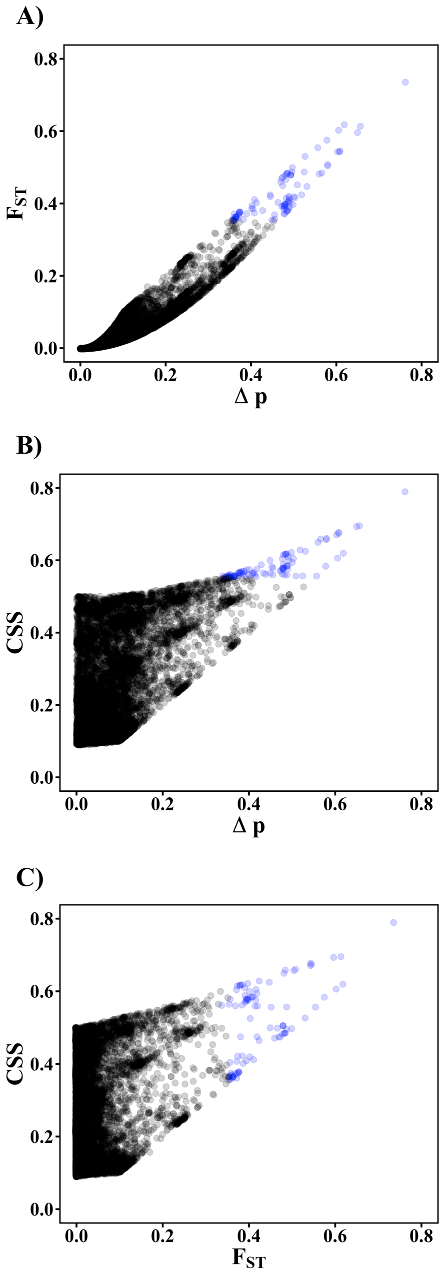

**Supplementary Methods S1. Phenotypic parallel evolution: parallel nucleotide site**

*Approach 1*: To detect outliers between the ecotypes (all Dune populations vs all Headland populations), we undertook three outlier detection methods: the top 1% from the distribution of $F_{ST}$ values, the top 1% from the distribution of cluster separation scores (CSS), and those SNPs identified by *BayeScan* (Foll and Gaggiotti 2008). $F_{ST}$ per SNP (Weir and Cockerham 1984) was calculated within *VCFtools* (Danecek et al. 2011), and we selected the top 1% (corresponding to $F_{ST} > 0.353$, which also corresponds to an average change in allele frequency, $\Delta p$, of 0.47, SD = 0.080; Figure S8A). We calculated the cluster separation score (CSS; Jones et al. 2012) per SNP using a custom R script (see GitHub [update-link-upon-acceptance] for details). CSS is a genetic distance-based measure that quantifies the average divergence between Dune and Headland clusters, after accounting for the variance within each ecotype. It ranges between 0 and 1, with higher values indicating a more distinct separation between Dune and Headland ecotypes. We selected the top 1% of CSS (corresponding to CSS > 0.523, an average $\Delta p$ of 0.45, SD = 0.087; Figure S8B). We also detected outliers in *BayeScan* v2.1 (Foll and Gaggiotti 2008). *BayeScan* implements a reversible-jump MCMC algorithm to estimate posterior odds comparing a model with and without selection for each SNP. We used default parameters with a prior odd of ten, meaning the neutral model is ten times more likely than the model with selection (as recommended for datasets of our size; Foll and Gaggiotti 2008). Results were robust to increasing the prior odds to 100 (data not shown). We categorized SNPs as highly differentiated if they contained a posterior probability > 0.91, corresponding to a Bayes Factor of > 10, and corresponding to an average $\Delta p$ of 0.38, SD = 0.121. For approach 1, we classified SNPs as outliers if they were detected in at least two of the three methods (Figure S8C).

*Approach 2*: We detected outliers separately for the Dune-Headland pairs at each locality. For the pair at each locality, SNPs for which MAF < 0.05 were filtered, retaining between 5,513 and 8,875 SNPs per pair (mean = 7,461 SNPs; SD = 1,038). As above, we identified outliers for each replicate pair using a combination of $F_{ST}$, CSS, and *BayeScan*. Instead of selecting the top 1% of $F_{ST}$ and CSS values (which is highly dependent on the number of sampled SNPs) we chose stringent cut-offs of $F_{ST}$ and CSS > 0.95. Within *BayeScan*, we considered SNPs with a posterior probability > 0.91 as highly differentiated, which corresponds to a Bayes Factor > 10. We considered a SNP an outlier per locality if it was detected as an outlier in at least two of the approaches ($F_{ST}$, CSS, and *BayeScan*). We considered a SNP an *Approach 2* outlier if it was detected as an outlier across all nine or eight replicate pairs.

*Approach 3*: To detect more subtle signals of outliers between the ecotypes, we asked whether there were concordant allele frequency changes across replicate pairs. Specifically, if a nucleotide site was highly differentiated in at least one pair according to *Approach 2*, we compared allele frequencies across all pairs for the site, and we asked whether the $\Delta p$ for each replicate pair was in the same direction across all localities (tested using two-sided dependent-samples sign-tests in R).

**Supplementary Methods S2. *Senecio lautus* transcriptome**

*Plant material*

We de-novo assembled a *S. lautus* transcriptome from seven different samples that came from different populations, tissues and plant developmental stages, using RNAseq (Wang et al. 2009). Specifically, we used plants at a young stage (before they flowered) to sample the shoot apical meristem (SAM), leaf primordia (LP) and the youngest three leaves (which were smaller than 500µm) from the main stem from 96 plants from each of four natural populations (see Table below for population details). We refer to these samples as SAM-LP samples. We used pools of plants because a single sample of this tissue does not yield enough material to carry out RNASeq (Ozsolak and Milos 2011). In addition, we also collected a fully expanded leaf smaller than 2cm, a fully expanded leaf bigger than 2cm and a flower from a single mature plant (i.e., after flowering) from H01 (see Table below).

*Growth conditions*

To obtain the SAM-LP tissue samples described above, seeds collected in the field from four natural populations (A03, D01, H01 and T01) were grown as follows: *S. lautus* seeds were cut at the shoot edge to speed-up and synchronize germination and placed in damped filter papers inside 50 mm diameter x 15 mm glass petri dishes. Seeds were kept in the dark for two days and then transferred to 12-hour day-night cycle for eight days in controlled temperature room at 25°C. Seedlings were then transferred to soil (70% pine bark, 30% coco peat, with 5kg/m$^3$ slow release Osmocote fertilizer) and were transferred to a greenhouse where they were watered as necessary. Plants were randomly crossed within each natural population to produce seeds. Next, 96 greenhouse produced seeds of each of the four natural populations were germinated following the protocol described above, but instead of transferring the seedlings to soil, seedlings were transferred to jiffy-7® peat pellets in 48-cell flats in a controlled temperature room at 25°C and a 12-hour day-night cycle. After seven weeks we dissected SAM, LP and the three youngest leaves from the main stem under a dissecting microscope. We pooled the 96 samples per population and stored leaf tissue in AMBION RNAlater®. In addition, a single plant from population H01 was grown as described above, and pruned after flowering to obtain abundant shoots. Approximately two weeks after pruning we collected a fully expanded leaf smaller than 2cm, a fully expanded leaf bigger than 2cm and a flower, which were snap-frozen in liquid nitrogen.

*RNASeq libraries*

Total RNA was extracted from each of the above seven samples using TRIzol® (Simms et al. 1993) and quantified using a NanoDrop (Thermo Fisher Scientific, Waltham, USA). Quality was checked using a bioanalyzer (Agilent technologies) at the Australian Genome Research Facility. We ensured samples contained an RNA integrity number (Schroeder et al. 2006) > 6.5 and a ratio of the 28S and 18S ribosomal bands > 1.0. For each sample 0.5ug of total RNA was used to construct sequencing libraries with the TruSeq RNA Sample Preparation Kits. Libraries were constructed and 100bp pair-end sequenced with Illumina HiSeq2000 at the Beijing Genomic Institute.

To develop a reference transcriptome for *S. lautus*, we first used SEECER v0.1.3 (Le et al. 2013) and default parameters to correct errors in RNAseq data. We then carried out de novo assembly using Trinity v2.0.2 (Grabherr et al. 2011). Following methods by Yang and Smith (2013), we chose the representative transcripts from each locus by retaining the transcript with the highest read coverage for each subcomponent. To further remove the redundant transcripts, which may come from alternative splicing or close paralogs, we clustered the assembly using CD-HIT-EST v4.6.1 (-c 96 -n 8 -r 1) and then chose 1 representative from each cluster.

**Supplementary Methods S2 Table. Plant samples for transcriptome assembly**

Details of the plants used to assemble the reference transcriptome. A03 coordinates: S36° 52.358' E147° 17.325'; T01 coordinates: S28° 13.831' E153° 8.105'. See Table S1 for coordinates for D01 and H01. SAM-LP refers to samples of the shoot apical meristem (SAM), leaf primordia (LP) and the youngest three leaves (which were smaller than 500µm). Young means plants before flowering and mature means plants after flowering. N represents the total number of plants used per sample.

| Population code | Ecotype | Tissue sampled | Plant stage | N |
|---|---|---|---|---|
| A03 | Alpine | SAM-LP | Young | 96 |
| T01 | Tableland | SAM-LP | Young | 96 |
| D01 | Dune | SAM-LP | Young | 96 |
| H01 | Headland | SAM-LP | Young | 96 |
| H01 | Headland | Expanded leaf < 2cm | Mature | 1 |
| H01 | Headland | Expanded leaf > 2cm | Mature | 1 |
| H01 | Headland | Flower | Mature | 1 |

**Supplementary Methods S3. Enrichment analysis**

For the enrichment analysis, sometimes there was more than one significant category within a cluster. To deal with this, we manually assigned an overall term for the cluster, based on the descriptions per category. See GitHub (update-link-upon-acceptance) for a step-by-step explanation of this process, including all enrichment output files and the summarized terms across replicate pairs.

**Supplementary Methods S4. Parallel auxin genes**

We examined whether there were auxin genes highly divergent across multiple localities within our current dataset. To do this, for each population pair we extracted any highly differentiated genes (as explained in the *Methods* section of the main text) that contained the word "auxin" in the gene name or gene ontology description. We then asked how many replicate pairs have each auxin gene as highly differentiated. These results are found in Table S19.

### References

Danecek, P., A. Auton, G. Abecasis, C. A. Albers, E. Banks, M. A. DePristo, R. E. Handsaker, G. Lunter, G. T. Marth, S. T. Sherry, G. McVean, R. Durbin, and 1000 Genomes Project Analysis Group. 2011. The variant call format and VCFtools. Bioinformatics 27:2156–2158.

Foll, M., and O. Gaggiotti. 2008. A genome-scan method to identify selected loci appropriate for both dominant and codominant markers: A Bayesian perspective. Genetics 180:977–993.

Grabherr, M. G., B. J. Haas, M. Yassour, J. Z. Levin, D. A. Thompson, I. Amit, X. Adiconis, L. Fan, R. Raychowdhury, Q. Zeng, Z. Chen, E. Mauceli, N. Hacohen, A. Gnirke, N. Rhind, F. di Palma, B. W. Birren, C. Nusbaum, K. Lindblad-Toh, N. Friedman, and A. Regev. 2011. Trinity: reconstructing a full-length transcriptome without a genome from RNA-Seq data. Nat. Biotechnol. 29:644–652.

James, M. E., H. Arenas-Castro, J. S. Groh, S. L. Allen, J. Engelstädter, and D. Ortiz-Barrientos. 2021. Highly replicated evolution of parapatric ecotypes. Accepted Mol. Biol. Evol., doi: 10.1101/2020.02.05.936401.

Jones, F. C., Y. F. Chan, P. Russell, E. Mauceli, J. Johnson, R. Swofford, M. Pirun, M. C. Zody, S. White, E. Birney, S. Searle, J. Schmutz, J. Grimwood, M. C. Dickson, R. M. Myers, C. T. Miller, B. R. Summers, A. K. Knecht, S. D. Brady, H. Zhang, A. A. Pollen, T. Howes, C. Amemiya, E. S. Lander, F. Di Palma, K. Lindblad-Toh, and D. M. Kingsley. 2012. The genomic basis of adaptive evolution in threespine sticklebacks. Nature 484:55–61.

Le, H.-S., M. H. Schulz, B. M. McCauley, V. F. Hinman, and Z. Bar-Joseph. 2013. Probabilistic error correction for RNA sequencing. Nucleic Acids Res. 41:e109.

Ozsolak, F., and P. M. Milos. 2011. RNA sequencing: advances, challenges and opportunities. Nat. Rev. Genet. 12:87–98.

Roda, F., H. Liu, M. J. Wilkinson, G. M. Walter, M. E. James, D. M. Bernal, M. C. Melo, A. Lowe, L. H. Rieseberg, P. Prentis, and D. Ortiz-Barrientos. 2013. Convergence and divergence during the adaptation to similar environments by an Australian groundsel. Evolution 67:2515–2529.

Schroeder, A., O. Mueller, S. Stocker, R. Salowsky, M. Leiber, M. Gassmann, S. Lightfoot, W. Menzel, M. Granzow, and T. Ragg. 2006. The RIN: an RNA integrity number for assigning integrity values to RNA measurements. BMC Mol. Biol. 7:3.

Simms, D., P. E. Cizdziel, and P. Chomczynski. 1993. TRIzol: A new reagent for optimal single-step isolation of RNA. Focus 15:532–535.

Wang, Z., M. Gerstein, and M. Snyder. 2009. RNA-Seq: a revolutionary tool for transcriptomics. Nat. Rev. Genet. 10:57–63.

Weir, B. S., and C. C. Cockerham. 1984. Estimating F-Statistics for the analysis of population structure. Evolution 38:1358.

Yang, Y., and S. A. Smith. 2013. Optimizing de novo assembly of short-read RNA-seq data for phylogenomics. BMC Genomics 14:328.
